# Generation of iPSCs from endangered Grevy’s zebra and comparative transcriptomic analysis of mammalian PSCs

**DOI:** 10.1101/2021.08.10.455807

**Authors:** Yoshinori Endo, Ken-ichiro Kamei, Koichi Hasegawa, Keisuke Okita, Hideyuki Ito, Shiho Terada, Miho Inoue-Murayama

**Author notes:** Corresponding authors Ken-ichiro Kamei, Institute for Integrated Cell-Material Sciences (WPI-iCeMS), Kyoto University, Kyoto, Japan, Miho Inoue-Murayama, Wildlife Research Center, Kyoto University, Kyoto, Japan.

## Abstract

Induced pluripotent stem cells (iPSCs) can provide a biological resource for functional and conservation research in various species. This expectation has led to generation of iPSCs from various species, including those identified as endangered species. However, the understanding of species variation in mammalian iPSCs is largely unknown. Here, to gain insight into the species variation in iPSCs, we the first generated iPSCs from the endangered species Grevy’s zebra (*Equus grevyi*; gz-iPSCs) for the first time in the world. We isolated primary fibroblasts cell from an individual that had died of natural causes at a zoo and reprogrammed the fibroblasts into iPSCs. We confirmed their pluripotency and differentiation potential and performed RNA sequencing analysis. The gz-iPSC transcriptome showed that the generated gz-iPSCs robustly expressed genes associated with pluripotency and reprogramming processes, including epithelial-to-mesenchymal and mesenchymal-to-epithelial transitions. Comparative transcriptomics with other species revealed patterns of gene expression among mammalian PSCs and detected evolutionary conservation of pluripotency-associated genes and the plausible importance of the translation process. This study provides new insights into the evolution of mammalian PSCs, and the species conservation and variation of PSCs will advance our understanding of the early development of mammals.

## Introduction

Mammalian induced pluripotent stem cells (iPSCs), which show unlimited self-renewal and differentiation capabilities into all three germ layers, can be potential sources of differentiated tissue cells for fundamental research and conservation of diverse species, especially those classified as endangered species. In general, biological materials of non-model mammals are constrained because of ethical and technical concerns, and the potential properties of PSCs enable the provision of resources for functional study and assisted reproductive technologies^1^. The development of iPSC technology^2, 3^ has broadened the opportunity to study PSCs from a range of mammalian species. Given, however, that even human and mice PSCs show different characteristics and the foundation of PSCs is yet to be completely elucidated^4^, it is of profound importance to understand the species variation and evolution of mammalian PSCs.

PSCs exhibit both similarities and differences in their characteristics between species, highlighting the importance of understanding PSCs from various species. Derivations of PSCs from a range of species have been reported, including cow^5^, pig^6^, horse^7–11^, naked mole-rat^12^, and other mammalian species^13, 14^. In the reprogramming of iPSCs, the defined combination of transcription factors can be effective with a wide range of taxonomic groups, except for some species that may require alternative factors, including bats, Tasmanian devils, platypus, and felids^15–19^. In most studies, PSCs have been shown to satisfy many of the criteria for pluripotency, while the characteristics of the cells are not completely defined^13, 14^. PSCs can reside in various pluripotency states, such as naive and primed pluripotency, and differences in pluripotency states and configurations have been reported between humans and mice^20^; besides, other species may exhibit alternative pluripotency states^13^. While biological processes and associated genes that take crucial roles in PSCs have been extensively studied in humans and mice, the molecular basis underlying the variation in mammalian PSCs is poorly explored.

The comparative genetic approach is a powerful tool for elucidating evolution^21^. While we previously described the evolutionary pattern in the pluripotency gene regulatory network from changes in protein-coding genes^22^, changes in gene expression may enable further insights into the phenotypic differences and similarities between species^23^. Comparative PSC gene expression analysis has previously highlighted the common regulation of signalling pathways between primates and mice^24^; evolutionary patterns across broader taxonomic lineages are poorly explored.

Compared to other taxonomic groups, Perissodactyla PSCs are exclusively limited in horse^7–11^, except for the Northern white rhinoceros^25, 26^. Grevy’s zebra (*Equus grevyi*) is one of the three extant zebra species and is the largest living wild equid. Grevy’s zebra has experienced a serious population decline of 54% over the last 30 years, leaving approximately 2,600 individuals^27^ and has been classified as the CITES Appendix I and ‘Endangered’ in the IUCN Red List. Grevy’s zebra belongs to the family Equidae, the taxonomic group including horses, donkeys, and zebras; thus, the adaptation of assisted reproduction techniques might be possible. Concerning the low genetic diversity of this species^28^, iPSCs from Grevy’s zebra might aid conservation efforts.

Here, we report the first generation of iPSCs from Grevy’s zebra (gz-iPSCs). We reprogrammed zebra fibroblasts by transducing four transcription factors, *OCT3/4* (also known as *POU5F1*), *SOX2*, *KLF4*, and *c-MYC*, using retroviral vectors. gz-iPSCs exhibited primed-type morphology and could be maintained under primed-type culture conditions and expressed pluripotency markers. To understand the molecular basis of the generated gz-iPSCs, we performed RNA sequencing (RNA-seq). In addition, we compared the transcriptome of Grevy’s zebra and other mammalian species and found evolutionary conservation and variations in gene expression pattern among mammalian PSCs. This study provides insights into the variations in mammalian PSCs and contributes to the future conservation management of endangered species.

## Methods

### Primary culture of Grevy’s zebra fibroblast

This study was conducted in strict accordance with the guidelines for the ethics of animal research by Kyoto University and the Wildlife Research Center of Kyoto University (WRC-2021-0016A). The sampling and methods were approved by the Kyoto City Zoo and the Wildlife Research Center of Kyoto University. Skin tissue samples were obtained from a female Grevy’s zebra that had died of natural causes at the Kyoto City Zoo (Japan). Primary fibroblasts were established as previously described^29^. The sample was sterilised with 70% (v/v) ethanol and cut into 1–2 mm^3^ pieces and was cultured in Dulbecco’s modified Eagle medium (DMEM) (Sigma-Aldrich, Merck, Darmstadt, Germany) with 10% (v/v) foetal bovine serum (FBS) (CCB, Nichirei Bioscience, Tokyo, Japan), 100 U/mL penicillin/streptomycin (Fujifilm Wako Pure Chemical Corporation, Osaka, Japan), 2.5 μg/mL amphotericin B (Sigma-Aldrich), and 100 µM non-essential amino acids (NEAA) (Sigma-Aldrich) in a humidified incubator at 37 °C with 5% (v/v) CO_2_. Fibroblast cultures at passage 3 were cryopreserved by suspending cells in CELLBANKER 1 (Takara Bio, Shiga, Japan), slowly cooled to -80 °C using a Mr. Frosty Freezing Container (Thermo Fisher Scientific, Waltham, MA, United States) for at least 24 h, and subsequently transferred to the liquid nitrogen vapour.

### Cell culture

Grevy’s zebra fibroblasts (gz-fibroblasts) were cultured in DMEM supplemented with 10% (v/v) FBS, 100 units/mL penicillin/streptomycin, and 100 µM NEAA on gelatin-coated dishes. Primate ES Cell Medium (ReproCELL, Kanagawa, Japan) with 100 µM sodium butyrate (Fujifilm Wako) and 10 µM Rho-associated coiled-coil forming kinase (ROCK) inhibitor Y-27632 (Fujifilm Wako) were used as primary iPSC medium. 0.3 µM glycogen synthase kinase-3 (GSK-3) inhibitor CultureSureR CHIR99021 (Fujifilm Wako), 0.1 µM ATP-competitive inhibitor CultureSureR (Fujifilm Wako) (correctively called 2i), and 1,000 U/mL leukaemia inhibitory factor (LIF) (Millipore, Merck, Darmstadt, Germany), collectively called 2i/LIF, were used with primary iPSC medium. After colonies appeared, putative gz-iPSCs were cultured in mTeSR-1 (Stemcell Technologies, Vancouver, Canada)^30^ on Matrigel (Corning, Corning, NY, United States)-coated dishes. Mouse embryonic fibroblasts (MEFs) were isolated from embryonic day 13.5 embryos of C57BL/6-Slc mice. Mouse fibroblasts SNL76/7 were clonally derived from a Sandos-inbred 6-thioguanine-resistant, ouabain-resistant (STO) cell line and stably express a neomycin-resistant cassette and a leukaemia inhibitory factor expression construct (MSTO)^31^. MEF and MSTO were cultured in DMEM supplemented with 10% (v/v) FBS, 100 units/mL penicillin/streptomycin, and 100 µM NEAA on gelatin-coated dishes. Fibroblasts were passaged using trypsin-EDTA (0.25%) (Thermo Fisher Scientific). gz-iPSCs were passaged using TrypLE Express (Thermo Fisher Scientific) with the addition of ROCK inhibitor at 10 µM 24 h before and after passaging. The cells were cultured in a humidified incubator at 37 °C with 5% (v/v) CO_2_. Cellular samples were tested for mycoplasma infection using the MycoAlert Mycoplasma Detection Kit (Lonza, Basel, Switzerland) according to the manufacturer’s protocol. All clones were expanded until at least passage 10 and then cryopreserved by suspending cells in STEM-CELLBANKER (Zenoaq, Fukushima, Japan), slowly cooled to -80 °C using a Mr. Frosty Freezing Container for at least 24 h, and subsequently transferred to the liquid nitrogen vapour.

### Virus production and generation of iPSCs

Cellular reprogramming was conducted using retrovirus vectors, as previously described^2, 3^. Briefly, pMXs-based retroviral vectors were prepared using human *OCT3/4*, *SOX2*, *KLF4*, and *c-MYC*^2^. Plat-GP packaging cells were seeded at 3 × 10^5^ cells per well in a 12-well plate^32^. The next day, 0.5 µg retroviral vectors were independently introduced into Plat-GP cells using 2.25 µL of FuGENE 6 transfection reagent (Promega, Madison, WI, United States). In addition, pMXs-EGFP was used to investigate transfection efficiency. To overcome the species barrier, the retroviruses were packaged with the VSV.G envelope protein using 0.25 µg pMD2.G. After 24 h, the medium was replaced with 1 mL of DMEM containing 10% FBS. Grevy’s zebra fibroblasts were seeded at 5 × 10^4^ cells/well in a 12-well plate. The next day, virus-containing supernatants from these Plat-GP cultures were collected and filtered through a 0.45-µm cellulose acetate filter. Virus-containing supernatants were either collected or concentrated by mixing with quarter volumes of 5 × PEG-it Virus Precipitation Solution (System Biosciences, Palo Alto, CA, United States), followed by centrifugation according to the manufacturer’s protocol. The retroviral pellet was suspended in DMEM and supplemented with polybrene at a final concentration of 4 µg/mL. Grevy’s zebra fibroblasts were transduced with viruses by incubating in a virus/polybrene-containing medium for 24 h. The cells were trypsinized 3 days after transduction, and 2–8 × 10^3^ cells were re-seeded on 6-well plates, 60 mm or 100 mm dishes coated with Matrigel, on mitomycin C-treated MSTO, or MEF feeder layer. The culture medium was replaced the next day with a primary iPSC medium with or without 2i/LIF. The number of colonies was counted on Day 17. The formed colonies were mechanically passaged in 96-well plates and individually expanded by further passaging. Two independent reprogramming experiments were performed, and the reprogramming conditions tested in this study are summarised in Table 1.

**Table 1.** Conditions of reprogramming experiments and number of iPSC-like colonies.

| Ex. ID | Scale | Num. cells | Vector concentration | Layer | 2i/LIF<sup>c</sup> | Num. colony |
| --- | --- | --- | --- | --- | --- | --- |
| 1 | 6-well | 2.00E+03 | As collected | MEF<sup>a</sup> | - | 0 |
| 1 | 6-well | 2.00E+03 | As collected | MEF | + | 3 |
| 1 | 6-well | 2.00E+03 | As collected | MSTO<sup>b</sup> | - | 1 |
| 1 | 6-well | 2.00E+03 | As collected | MSTO | + | 5 |
| 1 | 60 mm | 4.00E+03 | As collected | Matrigel | - | 2 |
| 1 | 60 mm | 4.00E+03 | As collected | Matrigel | + | 3 |
| 1 | 6-well | 2.00E+03 | Concentrated | MEF | - | 0 |
| 1 | 6-well | 2.00E+03 | Concentrated | MEF | + | 1 |
| 1 | 6-well | 2.00E+03 | Concentrated | MSTO | - | 2 |
| 1 | 6-well | 2.00E+03 | Concentrated | MSTO | + | 4 |
| 1 | 100 mm | 5.00E+03 | Concentrated | MSTO | + | 1 |
| 1 | 60 mm | 4.00E+03 | Concentrated | Matrigel | - | 30 |
| 1 | 60 mm | 4.00E+03 | Concentrated | Matrigel | + | 15 |
| 2 | 60 mm | 4.00E+03 | Concentrated | Matrigel | - | 16 |
| 2 | 60 mm | 8.00E+03 | Concentrated | Matrigel | - | 32 |

<sup>a</sup>Mouse embryonic fibroblasts, <sup>b</sup>SNL-STO <sup>c</sup>CHIR99021, PD0325901, and leukaemia inhibitory factor (LIF).

### Karyotyping

Karyotyping of putative gz-iPSCs was performed by the Nihon Gene Research Laboratories Inc. (Miyagi, Japan). The karyotypes of 50 cells were analysed using G-band staining, and the number of cells was counted according to the number of chromosomes.

### Alkaline phosphatase staining

Alkaline phosphatase (AP) staining was performed using the AP Staining Kit, AP100R-1 (System Biosciences) according to the manufacturer’s instructions. The cells were fixed with 4% (w/v) paraformaldehyde (Nisshin EM, Tokyo, Japan)/Dulbecco’s phosphate-buffered saline (DPBS) (Thermo Fisher Scientific) for 5 min at 24–26 °C and rinsed with DPBS. The cells were then stained with a freshly prepared staining solution for 20 min in the dark.

### Immunocytochemistry

Fluorescence immunocytochemistry was performed for the following pluripotency markers: OCT3/4 and NANOG. The cells were cultured for 3 days on glass-bottom dishes, fixed with 4% (w/v) paraformaldehyde/DPBS for 20 min at approximately 24–26 °C, rinsed twice with DPBS, and permeabilized with 0.5% (v/v) Triton X-100/DPBS (MP Biomedicals, Santa Ana, CA, United States) overnight at 4 °C. The cells were blocked with DPBS containing 5% (w/v) normal goat serum (Vector Laboratories, Burlingame, CA, United States), 5% (w/v) normal donkey serum (Jackson ImmunoResearch, West Grove, PA, United States), 3% (w/v) bovine serum albumin (BSA) (Sigma-Aldrich), and 0.1% (v/v) Tween20 (Bio-Rad Laboratories, Hercules, CA, United States) overnight at 4 °C and incubated with primary antibody diluted in blocking buffer overnight at 4 °C. The cells were washed twice with 0.1% (v/v) Tween20/D-PBS and incubated with secondary antibodies diluted in blocking buffer for 1 h at approximately 24–26 °C. After washing twice with 0.1% (v/v) Tween20/D-PBS, the nuclei were counterstained with 300 nM 4LJ,6-diamidino-2-phenylindole (DAPI) (Fujifilm Wako). The following primary antibodies were used at the indicated dilutions: mouse anti-OCT-3/4 (C-10 clone, #sc5279, 1:100) (Santa Cruz Biotechnology, Dallas, TX, United States) and rabbit anti-Nanog (#4903, 1:400) (Cell Signalling Technology, Danvers, MA, United States). The secondary antibodies used were labelled with anti-mouse IgG Alexa-488 or anti-rabbit IgG Alexa-594 (#715-546-150, #711-586-152, 1:1,000) (Jackson ImmunoResearch). Three gz-iPSC lines were tested as biological replicates. Grevy’s zebra fibroblasts and human iPSCs (253G1)^33^ (Supplementary method) were used as negative and positive controls for pluripotent markers, respectively. The experimental negative controls were tested by staining samples with only secondary antibodies.

### Gene expression of pluripotency markers

Total RNA was isolated using the RNeasy Mini Kit 50 (Qiagen, Hilden, Germany), according to the manufacturer’s instructions. DNA was eliminated with RNase-Free DNase Set (Qiagen) in solution, followed by RNA clean-up. Complementary DNA (cDNA) was synthesised using PrimeScript RT Master Mix (Takara Bio). Quantitative reverse transcription-polymerase chain reaction (qRT-PCR) analysis was performed using TB Green Premix Ex Taq II (Tli RNaseH Plus) (Takara Bio) on a Thermal Cycler Dice Real Time System TP800 (Takara Bio). The cycling conditions for qRT-PCR were as follows: 95 °C for 30 s, followed by 40 amplification cycles (95 °C, 5 s; 58 °C, 30 s). The relative expression ratios of target genes were calculated using the comparative Ct method and the expression levels of β*-actin* as the reference gene. The primers were designed using equine genomes as a reference with Primer-BLAST^34^ because Grevy’s zebra genome assembly and annotation are lacking at this time. Primers were designed to react specifically with the equine gene but not with humans for *OCT3/4, SOX2,* and *KLF4* (e*OCT3/4, eSOX2,* and e*KLF4*). Primers were also designed to react with both equine and humans for *OCT3/4, KLF4*, and *NANOG* (*ehOCT3/4, ehKLF4*, and *ehNANOG*). The primers used in this study are listed in Supplementary Table 1. Expression of pluripotency markers was assessed using gz-iPSCs as test samples and gz-fibroblasts as the somatic control sample. Dependency on 2i/LIF^35, 36^ was assessed by comparing the expression levels of pluripotency markers between samples cultured with or without 2i/LIF for three passages. Three independent experiments with three gz-iPSC clones were performed using qRT-PCR.

### *In vitro* embryoid body (EB) formation

Colonies of putative gz-iPSCs were mechanically cut into small aggregates of cells, detached from the culture dish with pipetting, and allowed to grow in suspension on ultra-low attachment culture dishes (Corning) in mTeSR-1 culture medium. After one week, the medium was replaced with a differentiation medium, DMEM supplemented with 20% (v/v) FBS, 100 units/mL penicillin/streptomycin, and 100 µM NEAA. Two weeks after the DMEM culture, samples were harvested for total RNA extraction. The ability to form derivatives of the three germ layers was assessed by gene expression of ectoderm, mesoderm, and endoderm markers using qRT-PCR, as previously described.

### Nanopore RNA-seq

Total RNA from gz-iPSCs and fibroblasts, each at three alternative generations (n_PSC_ = 3, n_fib_ = 3), was extracted as previously described and quantified using a NanoDrop 1000 spectrophotometer (Thermo Fisher Scientific) and a Bioanalyzer 2100 (Agilent Technologies, Santa Clara, CA, United States). RNA (40 ng) was used for library preparation using Oxford Nanopore Technologies (ONT, Oxford, United Kingdom) long-read cDNA sequencing. cDNA was generated using the PCR-cDNA Barcoding kit (SQK-PCB109) of ONT according to the manufacturer’s protocol. For sequencing, libraries were applied to the Nanopore Flow Cell (v 9.4.1) and run for up to 72 h.

### RNA-seq analysis

Sequenced reads were base-called and demultiplexed using the ONT EPI2ME software. Adapter sequences were trimmed from the reads using Porechop (v. 0.2.4)^37^. Low-quality reads were filtered using a NanoFilt included in the NanoPack (v. 2.7.1)^38^. The filtered reads were mapped to the horse transcriptome using the Minimap2 (v. 2.17)^39^. The mapped reads were counted using Salmon (v. 1.3.0)^40^. The transcriptome data of cow (n_PSC_ = 2, n_fib_ = 2) (PRJNA432600)^5^, human (n_PSC_ = 4, n_fib_ = 2) (PRJNA230824)^41^, mouse (n_PSC_ = 4, n_fib_ = 1) (PRJNA564252)^42^, NMR (n_PSC_ = 4, n_fib_ = 2) (PRJDB4191)^12^, and pig (n_PSC_ = 2, n_fib_ = 1) (PRJDB5113)^6^ PSCs and fibroblasts were collected from the European Nucleotide Archive (ENA; https://www.ebi.ac.uk/ena)^43^ database. Because the transcriptome of horse PSCs was not available, the transcriptome of horse inner cell mass (ICM) (n_PSC_ = 3) (PRJNA223157)^44^ was used and treated as PSCs. The transcriptome data were sequenced using Illumina platforms. Illumina reads were trimmed and filtered using a fastp (v. 0.20.0)^45^. The filtered reads were mapped to the transcriptome of each species and counted using salmon. The transcript reads were converted to gene-level abundance using tximport (v. 3.13)^46^ and annotated with human orthologues using the Biomart tool of Ensembl (http://www.ensembl.org/biomart/martview/)^47^. Differentially expressed genes (DEGs) were identified using DESeq2 (v. 1.28.1)^48^ with an FDR-adjusted *P-*value of 0.1, and | log2FoldChange | > 1 as default^49^. A volcano plot was constructed using EnhancedVolcano (v. 1.6.0), in which log2FoldChnage values were shrunken using the Apeglm method in DESeq2^50^ and FDR lower than 10E-20 were compressed for visualisation. Protein analysis through evolutionary relationships (PANTHER) provided by the Gene Ontology Consortium (http://geneontology.org)^51–53^ was used for gene ontology (GO) analysis of biological processes enriched for DEGs with FDR < 0.05. For DEG analysis across species, we combined data from all study species, compared the changes in gene expression between cell types, and used the top 1,000 DEGs according to FDR for later analysis. Hierarchical clustering and heat maps were constructed across species using the heatmap2 in gplots R package (v. 3.1.1) with the rlog transformation in DESeq2. DEGs per species were identified using the top 1,000 DEGs across the species. Venn diagrams were constructed using the VennDiagram R package (v. 1.6.0). Gene set enrichment analysis (GSEA) with Kyoto Encyclopedia of Genes and Genomes (KEGG) pathways^54^ was performed using the clusterProfiler R package (v. 3.16.1)^55^.

### Statistics and reproducibility

The Welch two-sample t-test was conducted using R (v. 4.0.3). Box plots were constructed using Python graphing packages Matplotlib (v. 3.3.4)^56^ and Seaborn (v. 0.11.1)^57^. Centre lines indicate median and box limits indicate upper and lower quartiles. Upper whisker = min(max(*x*), Q_3 + 1.5 × IQR), lower whisker = max(min(*x*), Q_1 - 1.5 × IQR).

## Results

### Generation of Grevy’s zebra iPSCs from primary fibroblasts

To acquire the source for gz-iPSCs, we obtained primary fibroblasts from the skin tissue of an adult female Grevy’s zebra (Figs. 1a and b). gz-fibroblasts grew in a commonly used cell-culture medium, such as DMEM supplemented with 10% (v/v) FBS. We confirmed that the gz-fibroblasts propagated until passage 10.

**Fig. 1.**
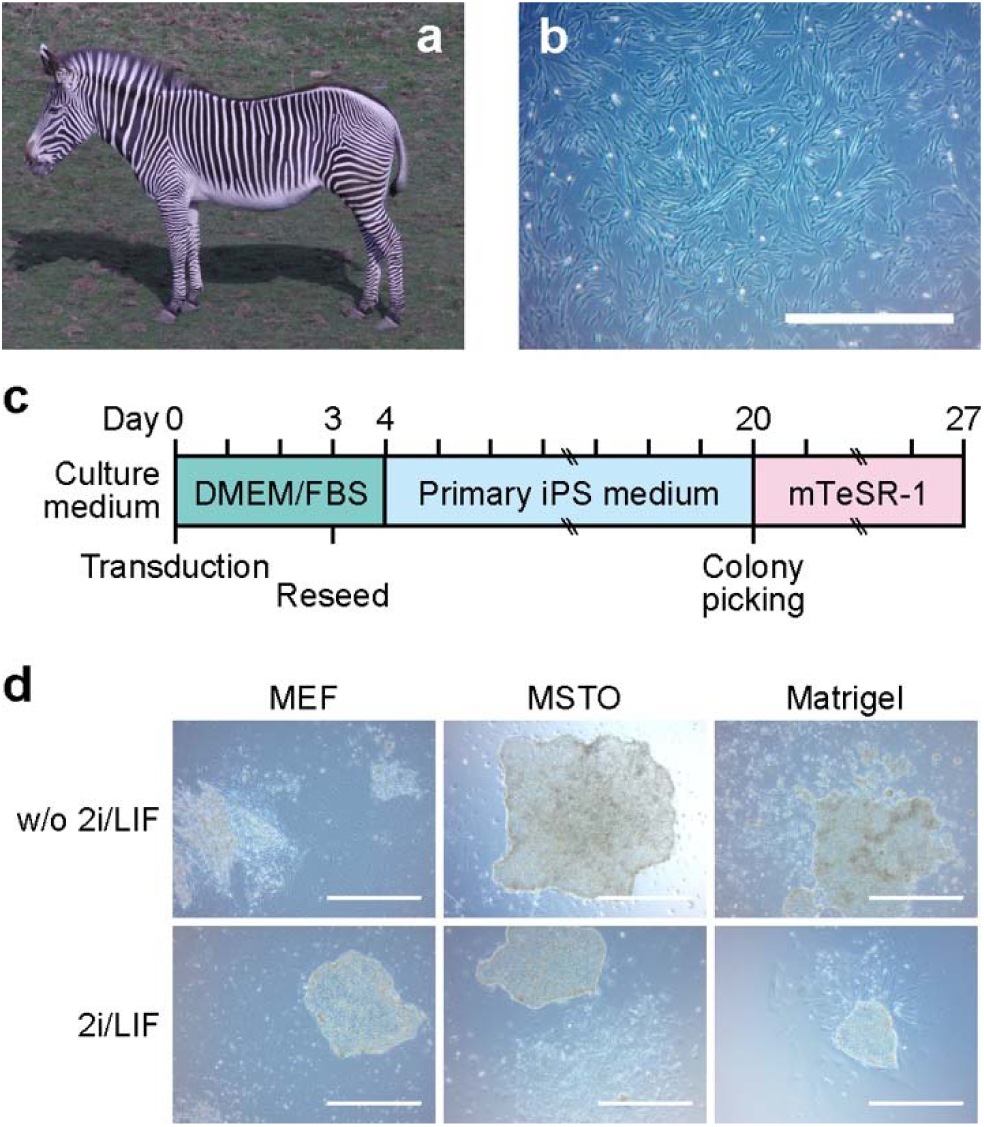
Generation of Grevy’s zebra iPSCs from primary fibroblasts. **a**, The Grevy’s zebra. **b**, Morphology of zebra fibroblasts. **c**, Schematic of the induction protocol. **d,** Morphology of induced colonies on day 17–20 of reprogramming procedure in different conditions. No PCS-like colony was observed in MEF without 2i/LIF condition. The scale bar represents 1,000 μm. iPSCs, induced pluripotent stem cells; DMEM, Dulbecco’s modified Eagle medium; FBS, fetal bovine serum; MEF, mouse embryonic fibroblasts; MSTO, mouse SNL-STO; 2i/LIF, CHIR99021, PD0325901 (2i), and leukaemia inhibitory factor (LIF).

To identify an efficient method for transgene delivery, we transduced gz-fibroblasts at passage three with retroviruses designed to express the human *OCT3/4*, *SOX2*, *KLF4*, and *c-MYC* (Fig. 1c) with an unconcentrated or concentrated vector, which has increased by viral titres and reduced toxicity^58^. gz-fibroblasts were resistant to viral toxicity, and the concentrated viral vectors exhibited higher transduction rates (Supplementary Fig. 1). To identify the efficient culture conditions for the formation of colonies, we reseeded transduced cells on Matrigel, which provides feeder-free surfaces for PSCs^59^, feeder layers, MEFs, and MSTOs, which secrete a variety of growth factors and extracellular matrices and are widely used in establishing PSC lines from a variety of species^2, 13, 60, 61^. Whereas our primary iPSC medium can sustain primed-type PSCs^2^, the formed colonies may exhibit naive-type characteristics, requiring distinct culture condition^35, 36^. To address this, we also tested the 2i/LIF condition in the cells after transduction until colony formation. PSC-like colonies formed on day 11 after transduction, followed by new colonies appearing periodically over the next 10 d (Supplementary Fig. 2). PSC-like colonies formed under all conditions, except in populations cultured with or without 2i/LIF on MEF, and the morphologies of the colonies were similar between conditions (Fig. 1d, Supplementary Fig. 3). Finally, we observed the highest number of colonies with a condition in which cells were transduced with a concentrated vector and cultured without 2i/LIF on Matrigel (Table 1).

### Characterisation of the pluripotent status of Grevy’s zebra iPSCs

To determine the culture conditions for maintaining gz-iPSCs, we compared cellular growth in the primary iPSC medium and the alternative mTeSR-1 medium. Given the high number of colonies observed in the Matrigel condition in our reprogramming experiment, we tested the mTeSR-1 medium, which has been developed for feeder-free culture of primed-type human PSCs^30^. The mTeSR-1 medium enabled the putative gz-iPSCs to grow stably, while the primary iPSC medium could not sustain colonies for more than a few passages (Supplementary Fig. 4). To determine whether gz-iPSCs can grow in naive-type condition^35, 36^, we cultured the gz-iPSCs with 2i/LIF in mTeSR-1 medium and observed a decrease in pluripotency markers in the presence of 2i/LIF (Supplementary Fig. 5). Therefore, we chose mTeSR-1 medium without 2i/LIF as the maintenance culture condition. We initially selected a total of 48 colonies from the most efficient condition in the reprogramming process, five of which could be maintained for up to at least five passages. To select the primary clones for continuous culture and later analyses, we performed preliminary pluripotency experiments. We selected three primary clones (named A, D, and E) based on the AP activity and the expression of pluripotency marker genes and the silencing of viral genes (Supplementary Fig. 6).

To determine whether the generated colonies exhibit the nature of mammalian PSCs, we investigated the cellular characteristics of the putative gz-iPSCs. The morphology of the gz-iPSCs resembled primed-type PSC colonies generated from humans, such as a monolayer of cells with clear colony edges, rather than mouse iPSCs, such as a semi-spherical colony (Fig. 2a). Nevertheless, the gz-iPSCs exhibited abundant cytoplasm compared to large nuclei and scant cytoplasm in human and mouse iPSCs^2, 3^. In addition, the colonies of gz-iPSC showed lose and sharp edges compared to that of human and mouse in which cell-cell tight junctions form round edges. The gz-iPSCs could be passaged as single cells with TrypLE Express even without ROCK inhibitor, which is required for survival of dissociated human ESCs^62^, while ROCK inhibitor improved the survival of gz-iPSCs (Supplementary Fig. 7). During passage from 24 to 28, gz-iPSCs showed doubling times of 22.6 ± 2.4 h, which is similar to that of human ESCs^63^. We investigated the chromosomal complement of the clone A and found that gz-iPSCs (68%) had a normal karyotype at passage 13, while 32% of them had one extra chromosome (Fig. 2b). The MycoAlert test on the supernatant of the gz-iPSCs showed that all three clones were negative for mycoplasma (Supplementary Fig. 8). To date, these gz-iPSC lines have been maintained for more than 30 passages.

**Fig. 2.**
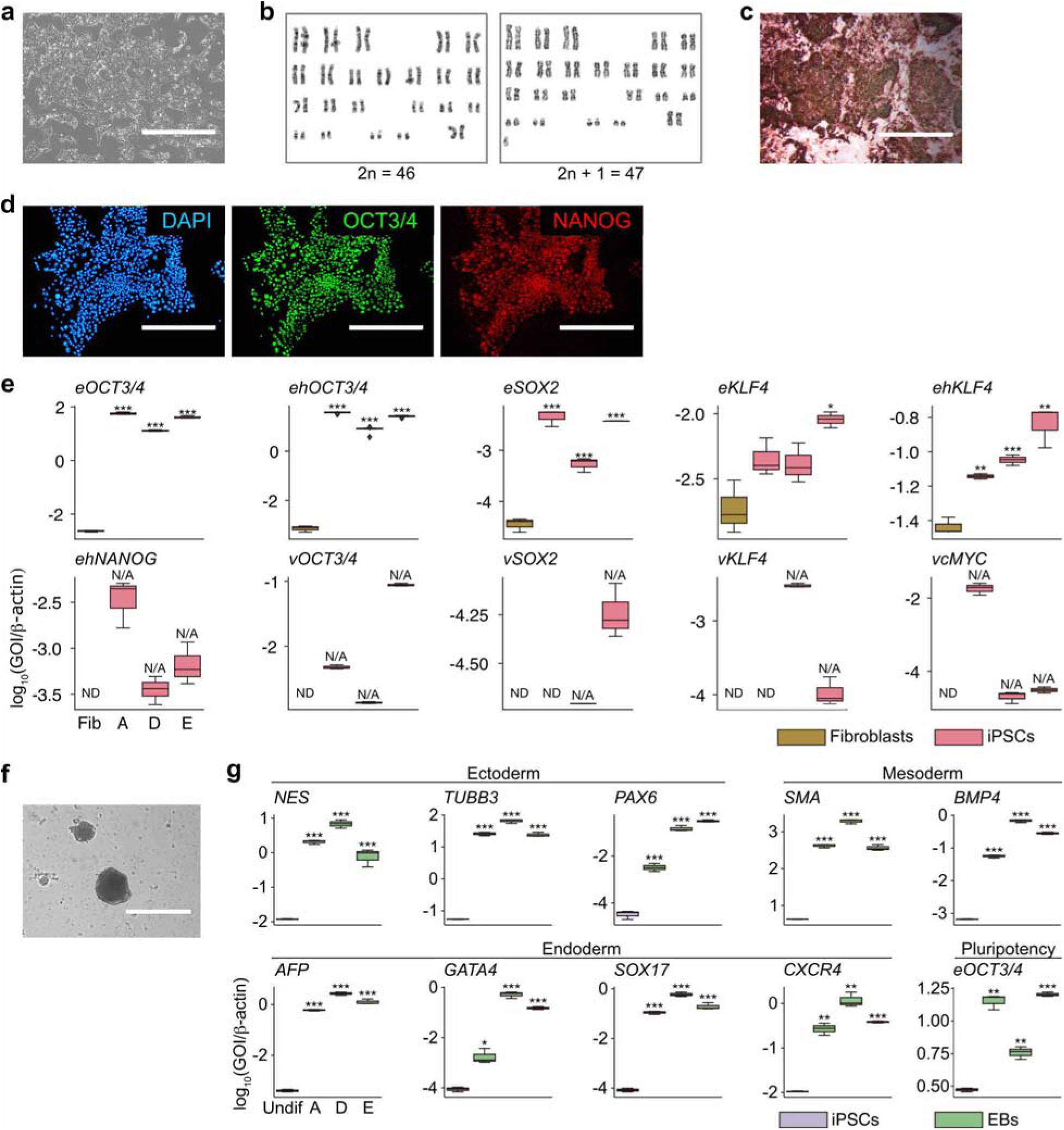
Characterisation of the pluripotent status of Grevy’s zebra iPSCs. **a**, Morphology of gz-iPSCs at passage 32. **b**, Karyotype. **c**, Alkali phosphatase activity. Clone D at passage 8 is shown here. **d**, Immunofluorescence for pluripotency markers. Nuclei are stained with DAPI. Clone A at passage 17 is shown here. **e**, **f**, and **h**, Box plots showing qRT-PCR. **e**, Comparison of pluripotency and viral marker expressions between gz-iPSCs at passage 25 and fibroblasts. **f**, Comparison of pluripotency marker expressions between gz-iPSCs cultured with or without 2i/LIF. **g**, Morphology of embryoid bodies after 7 days of differentiation in suspension culture. **h**, Comparison of three germ layer marker expressions between EBs and gz-iPSCs after two weeks of suspension culture in differentiation medium. A, D, and E represents independent clones of gz-iPSCs. Three independent experiments with three lines of iPSCs sample and fibroblasts (**e**) and three lines of EBs and undifferentiated gz-iPSC clone A (**h**) are shown here. Centre lines indicate median and box limits indicate upper and lower quartiles. Upper whisker = min(max(*x*), Q_3 + 1.5 × IQR), lower whisker = max(min(*x*), Q_1 - 1.5 × IQR). Statistical analyses were performed using the Welch two-sample t-test (* *P* < 0.05, ** *P* < 0.01, and *** *P* < 0.001). (**e**), *eOCT3/4*, A-Fib, df = 3.8059, *P* = 4.97 × 10^-8^, D-Fib, df = 3.9563, *P* = 2.6 × 10^-8^, E-Fib, df = 3.5094, *P* = 2.5 × 10^-7^; *ehOCT3/4*, A-Fib, df = 2.1301, *P* = 0.0001701, D-Fib, df = 4.6057, *P* = 4.16 × 10^-7^, E-Fib, df = 2.2128, *P* = 0.0001407; *eSOX2*, A-Fib, df = 3.8859, *P* = 8.24 × 10^-5^, D-Fib, df = 3.9932, *P* = 0.0004939, E-Fib, df = 2.0059, *P* = 0.001494; *eKLF4*, A-Fib, df = 3.6083, *P* = 0.06292, D-Fib, df = 3.703, *P* = 0.0842, E-Fib, df = 2.3481, *P* = 0.02099; *ehKLF4*, A-Fib, df = 2.3778, *P* = 0.005432, D-Fib, df = 3.3346, *P* = 0.0007948, E-Fib, df = 2.6524, *P* = 0.006234. (**g**), *NES*, A-Undif, df = 2.1631, *P* = 0.0001675, D-Undif, df = 2.0486, *P* = 0.0005409, E-Undif, df = 2.0101, *P* = 0.006796; *TUBB3*, A-Undif, df = 2.1178, *P* = 8.5 × 10^-5^, D-Undif, df = 2.0881, *P* = 9.55 × 10^-5^, E-Undif, df = 2.0758, *P* = 0.0001593; *PAX6*, A-Undif, df = 3.9738, *P* = 0.0001727, D-Undif, df = 3.4486, *P* = 3.26 × 10^-5^, E-Undif, df = 2.2265, *P* = 0.0004007; *SMA*, A-Undif, df = 2.1033, *P* = 0.000147, D-Undif, df = 2.0725, *P* = 0.0001293, E-Undif, df = 2.0387, *P* = 0.0005248; *BMP4*, A-Undif, df = 2.249, *P* = 8.19 × 10^-5^, D-Undif, df = 2.2438, *P* = 3.13 × 10^-5^, E-Undif, df = 2.466, *P* = 8.63 × 10^-6^; *AFP*, A-Undif, df = 3.4689, *P* = 4.86 × 10^-7^, D-Undif, df = 3.6565, *P* = 4.51 × 10^-7^, E-Undif, df = 2.9091, *P* = 1.76 × 10^-5^; *GATA4*, A-Undif, df = 2.3655, *P* = 0.01166, D-Undif, df = 3.4792, *P* = 9.19 × 10^-6^, E-Undif, df = 3.5874, *P* = 6.68 × 10^-7^; *SOX17*, A-Undif, df = 3.9355, *P* = 6.68 × 10^-7^, D-Undif, df = 3.7523, *P* = 1.15 × 10^-6^, E-Undif, df = 2.9946, *P* = 4.14 × 10^-5^; *CXCR4*, A-Undif, df = 2.028, *P* = 0.002962, D-Undif, df = 2.0191, *P* = 0.002084, E-Undif, df = 2.946, *P* = 2.26 × 10^-6^; *eOCT3/4*, A-Undif, df = 2.23, *P* = 0.001477, D-Undif, df = 2.3194, *P* = 0.006164, E-Undif, df = 3.8573, *P* = 7.91 × 10^-7^. In all statistical tests, sample size is n = 3, except *ehOCT3/4* with A, D, and E, n = 6. Scale bar represents 400 μm in **d** and **g**. 1,000 μm in **a** and **c**. iPSCs, induced pluripotent stem cells; Fib, fibroblasts; Undif, undifferentiated gz-iPSC clone A; DAPI, 4LJ,6-diamidino-2-phenylindole; 2i/LIF, CHIR99021, PD0325901 (2i), and leukaemia inhibitory factor (LIF); EBs, embryoid bodies; ND, not detected; N/A, not analysed; df, degrees of freedom; *e*, *h*, and *v* represent genes of equine, human, and virus, respectively.

To evaluate the expression of proteins associated with pluripotency, we conducted molecular staining followed by microscopic observation. In one of the pluripotent-associated proteins, the level of AP was observed after treatment with red-coloured substrates that reacted with AP at passages 6, 8, and 10 for clones A, D, and E, respectively (Fig. 2c and Supplementary Fig. 6a). Moreover, immunocytochemistry revealed the expression of pluripotency marker proteins (OCT3/4 and NANOG) with gz-iPSCs at passage 17 (Fig. 2d). We observed no fluorescent expression in the fibroblasts and the negative controls (Supplementary Fig. 9). The fluorescent expression of NANOG, which had not been transduced by a retroviral vector, supports the increase of the pluripotency marker protein in the reprogrammed gz-iPSCs.

For further evaluation of pluripotency criteria, we analysed the gene expression of pluripotency markers in gz-iPSCs using qRT-PCR. To determine whether the expressed genes were endo-or exogenous, we designed equine-specific primers, *eOCT3/4*, *eSOX2*, and *eKLF4*. We also designed multi-species-specific primers to react with both equine and human genes, named *ehOCT3/4*, *ehKLF4*, and *ehNANOG*, to confirm the expression of these markers. We observed higher expression levels of all the analysed pluripotency markers with iPSC samples compared with fibroblasts with both equine-specific and multi-species-specific primers at passage 25 (Fig. 2e). The expression of virally transduced genes (*vOCT3/4*, *vSOX2*, *vKLF4,* and *vcMYC*) was not completely silenced and was observed at low levels. However, the expression levels of the endogenous genes (*eOCT4*, *eSOX2*, and *eKLF4*) were much higher than those of the exogenous genes. Additionally, the generated gz-iPSCs expressed *NANOG*, which was not introduced in the reprogramming process and not observed in the original gz-fibroblasts. These results indicate that endogenous pluripotency genes were induced by the reprogramming process and maintained the generated gz-iPSCs.

To examine the differentiation ability of gz-iPSCs, we conducted EB formation, in which cells of all three germ layers were mixed (Fig. 2f). As observed in human iPSCs, gz-iPSCs formed ball-like EBs in suspension culture for two weeks with a differentiation medium. qRT-PCR analysis revealed increased expression of lineage markers for the three germ layers, including ectoderm (*NES*, *TUBB3*, and *PAX6*), mesoderm (*SMA* and *BMP4*), and endoderm (*AFP*, *GATA4*, *SOX17*, and *CXCR4*)^29^ (Fig. 2g and Supplementary Table 1).

### Identification of genes altered by the generation of gz-iPSCs

To investigate the comprehensive changes in gene expression by reprogramming, we performed RNA-seq and analysed the DEGs between gz-iPSCs and fibroblasts. DEG analysis revealed 1,144 upregulated and 1,495 downregulated DEGs with adjusted *P-*values (false discovery rate [FDR] < 0.1) and | log2FoldChange | > 1 by RNA-seq (Fig. 3a, Supplementary Table 2). As expected, the upregulated genes included the well-known pluripotency genes highly expressed compared to fibroblasts, such as *OCT3/4*, *DNMT3B*, *SALL4*, *ZFP42* (also known as *REX1*), and *LIN28*^64^. In contrast, the downregulated genes included fibroblast genes *VIM*, *DDR2*, *TGFBR2, COL1A1, COL1A2,* and *FSP1* (also known as *S100A4*)^65^.

**Fig. 3.**
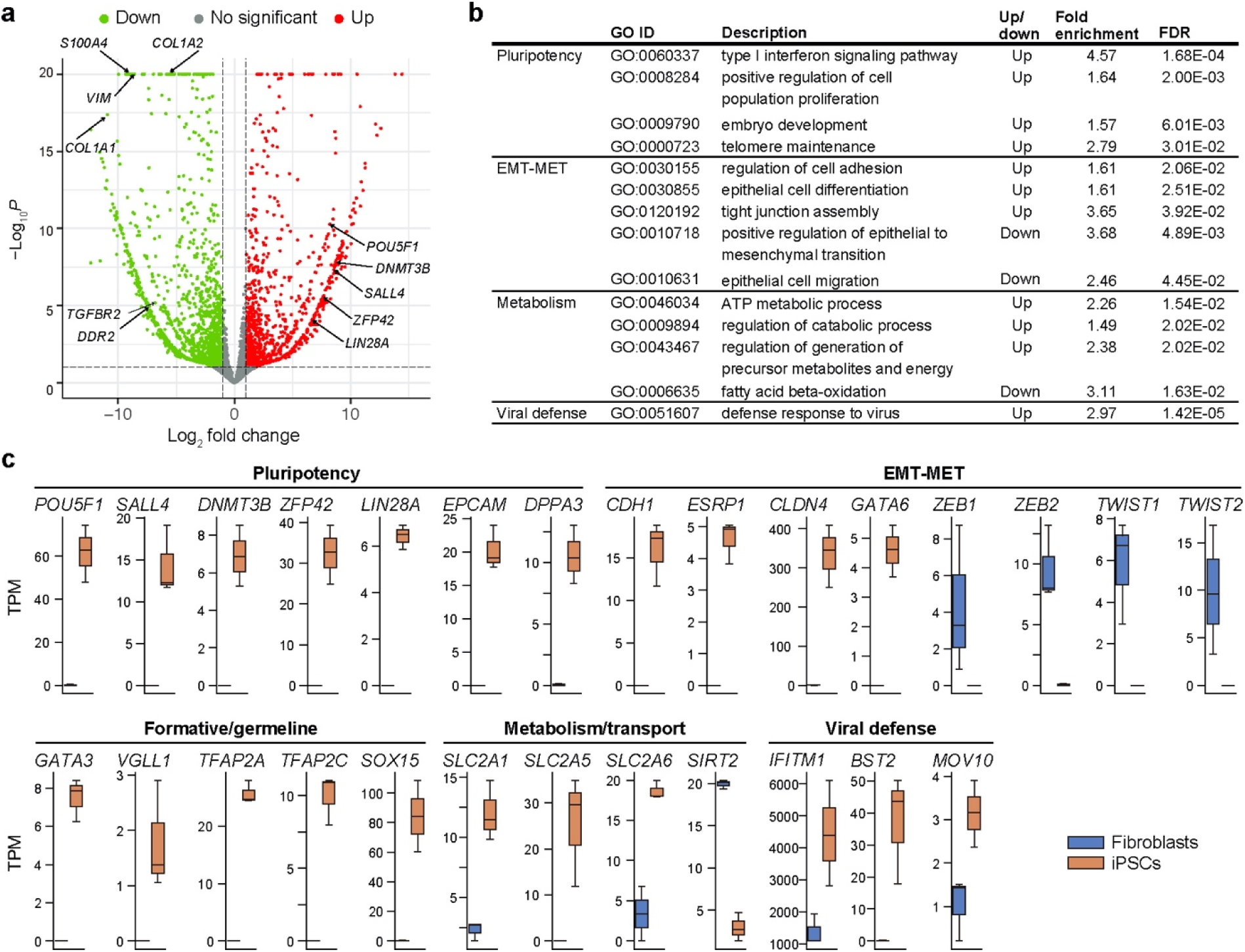
Differentially expressed gene (DEG) analysis of Grevy’s zebra fibroblasts and iPSCs transcriptome. **a,** Volcano plot showing the significant DEGs. The coloured dot represents gene that is upregulated (*red*) and downregulated (*green*) in gz-iPSCs (FDR < 0.1, | log2FoldChange | > 1). Log2FoldChange values were shrunken with the apeglm method and FDR lower than 10E-20 were compressed for visualisation. **b**, GO terms that are enriched with DEGs upregulated or downregulated in gz-iPSCs (FDR < 0.05). **c**, Box plots showing TPM of individual DEGs. All DEGs shown here are significantly different between iPSC and fibroblasts. Centre lines indicate median and box limits indicate upper and lower quartiles. Upper whisker = min(max(*x*), Q_3 + 1.5 × IQR), lower whisker = max(min(*x*), Q_1 - 1.5 × IQR). gz-iPSCs, Grevy’s zebra induced PSCs; DEG, differentially expressed gene; GO, gene ontology; FDR, false discovery rate; EMT-MET, epithelial-to-mesenchymal and mesenchymal-to-epithelial transitions; TPM, transcripts per million.

To characterise the derived gz-iPSCs for representative biological functions, we performed GO enrichment analysis (Fig. 3b and Supplementary Tables 3 and 4). Among the hierarchically specific subclasses, the GO terms enriched with upregulated DEGs included the type I interferon signalling pathway, positive regulation of cell population proliferation, embryo development, and telomere maintenance. GO analysis also revealed enrichment of the terms related to epithelial-to-mesenchymal and mesenchymal-to-epithelial transitions (EMT-MET). The upregulated EMT-MET terms included regulation of cell adhesion, epithelial cell differentiation, and tight junction assembly, whereas the downregulated terms included positive regulation of EMT, and epithelial cell migration. GO terms associated with metabolism included ATP metabolic process, regulation of catabolic process, and regulation of generation of precursor metabolites and energy with upregulated DEGs, as well as fatty acid beta-oxidation with downregulated DEGs.

To gain insights into the molecular basis underlying the enriched biological processes, we compared the transcripts per million (TPM) of the DEGs between gz-iPSCs and fibroblasts (Fig. 3c). In addition to the pluripotency signature genes shown in the volcano plot, we found upregulation of *EPCAM* and *DPPA3*. Among the EMT-MET-related biological processes, we observed upregulation of *CDH1* (also known as E-cadherin), which promotes MET^66^, *ESRP1*, which promotes MET via the upregulation of *CDH1*^67^, *CLDN4*^68^ and *GATA6*^69^, which suppress EMT. We also observed the downregulation of *ZEB1*, *ZEB2*, *TWIST1*, and *TWIST2*, that are highly expressed in EMT and suppressed in MET^70^. We also found upregulation of metabolic and glucose transport-associated genes *SLC2A1*, *SLC2A5*, and *SLC2A6* (*GLUT1*, *5*, and *6*, respectively) and downregulation of *SIRT2*. Furthermore, we found upregulation of *IFITM1*, *BST2* (*CD317*), and *MOV10*, which are involved in viral defence. To further evaluate the expression changes in DEGs, we inspected log2FoldChanges and found EMT-MET-related genes among the highly upregulated DEGs with log2FoldChange > 4, including *DMKN*, which is the key regulator of EMT^71^, *CDH1*, *CLDN4*, *EPCAM*, *ESRP1*, and *GATA6* (Supplementary Table 2). In addition, highly upregulated DEGs included markers of pluripotency state and germline cells, including *GATA3*, *VGLL1*, *TFAP2A*, *TFAP2C*, and *SOX15*.

### Gene expression pattern of mammalian PSCs

Comparative analysis of gene expression provides insights into the evolution of molecular basis among species^23^. To understand gz-iPSCs from an evolutionary perspective, we compared the transcriptome of Grevy’s zebra with other mammalian species, including human^41^, mouse^42^, naked mole-rat (NMR)^12^, cow^5^, and pig^6^, for whom the transcriptomes of PSCs and fibroblasts were available in public databases. As a reference for equine PSCs, we included the transcriptome of horse ICM^44^ because no transcriptomic data of horse PSCs were available. We excluded genes that were found in human samples only, putatively due to annotation bias, as these genes may cause clustering problems (Supplementary Fig. 10). To investigate the patterns of gene expression, we identified DEGs between fibroblasts and PSCs across species (Supplementary Table 5). Hierarchical clustering separated samples by cell type rather than by species (Fig. 4a). A visual inspection of the z-score shows that PSCs exhibited a higher degree of dispersion between species than that between fibroblasts. The hierarchical pattern does not reflect known phylogeny in both PSCs and fibroblasts, except for pairs of Grevy’s zebra-horse and cow-pig. The clade of PSCs contains two major groups, one of which includes Grevy’s zebra, most closely grouped with horse next with NMR, and the other includes cows, pigs, humans, and mice.

**Fig. 4.**
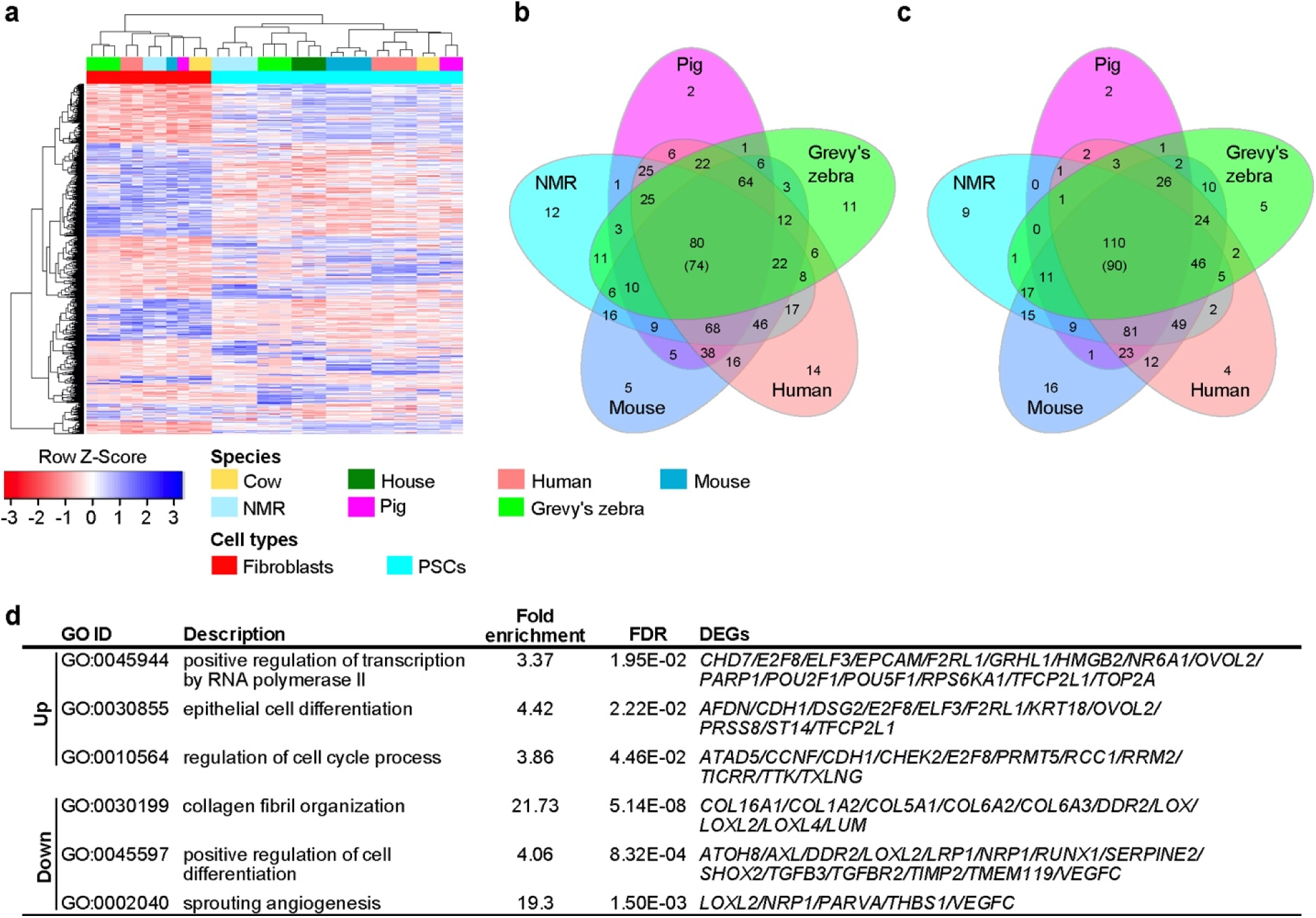
Gene expression pattern of mammalian PSCs. **a**, Heat map with hierarchical clustering of DEGs between PSCs and fibroblasts across species. The colour bars at the top indicate species, and the bars at the bottom indicate cell types. **b** and **c**, Venn diagram showing unique and common DEGs per species. **b,** Upregulated. **c**, Downregulated. Because it is difficult to include six or more elements in the Venn diagram, cow is excluded here. The number in brackets represents gene number including cow. **d**, GO terms that are significantly enriched with commonly upregulated or downregulated DEGs with FDR < 0.05. GO terms representative for PSC characteristics are shown here. Top 1,000 DEGs based on FDR are used here. PSCs, pluripotent stem cells; DEGs, differentially expressed genes; NMR, naked mole-rat; GO, gene ontology; FDR, false discovery rate.

To investigate the species differences in expression changes, we identified DEGs per species using the top 1,000 significant DEGs across mammals. DEG analysis revealed 74 commonly upregulated and 90 downregulated genes across the species (Fig. 4b and c). Among the DEGs with the lowest *P-*values, the commonly upregulated genes included well-known pluripotency-associated genes in both humans and mice, such as *ESRP1*, *EPCAM*, *OCT3/4*, *DNMT3B*, and *DSG2* (Table 2 and Supplementary Table 6). In addition, we found genes that are not generally associated with pluripotency, including *AP1M2*, *PLEKHA7*, and *MARVELD3*. The commonly downregulated DEGs include *S100A4*, which is a typical fibroblast marker^65^, and *DCN*, which inhibits ESC self-renewal^72^. GO terms enriched with upregulated common DEGs included positive regulation of transcription by RNA polymerase II, epithelial cell differentiation, and regulation of cell cycle process (Fig. 4d). The downregulated GO terms included collagen fibril organisation, positive regulation of cell differentiation, and sprouting angiogenesis (Supplementary Tables 7 and 8).

**Table 2.** List of top 20 DEGs^a^ across mammalian PSCs^b^ based on FDR^c^ by DESeq2. The expression changes per species are also shown.

| Gene | FDR | Cow | Human | Mouse | NMR<sup>d</sup> | Pig | Grevy's zebra |
| --- | --- | --- | --- | --- | --- | --- | --- |
| *ESRP1* | 2.19E-51 | + | + | + | + | + | + |
| *S100A4* | 1.28E-46 | - | - | - | - | - | - |
| *EPCAM* | 8.23E-37 | + | + | + | + | + | + |
| *DCN* | 3.15E-36 | - | - | - | - | - | - |
| *RFTN2* | 3.97E-34 | - | - | - | - | - | - |
| *AP1M2* | 2.43E-32 | + | + | + | + | + | + |
| *FAP* | 2.43E-32 | - | - | - | - | - | - |
| *COL6A3* | 1.11E-30 | - | - | - | - | - | - |
| *SRPX2* | 3.60E-27 | - | - | - | - | - | - |
| *GRB7* | 1.06E-26 | + | + | + | + | Nd<sup>e</sup> | + |
| *PRRX1* | 3.45E-26 | - | - | - | - | - | - |
| *CEMIP* | 3.45E-26 | - | - | - | ND | - | - |
| *CTSK* | 1.57E-25 | - | - | - | - | NR<sup>f</sup> | - |
| *POU5F1* | 1.57E-25 | + | + | + | + | + | + |
| *MAP1A* | 3.45E-25 | - | - | - | - | - | - |
| *EHD2* | 6.00E-25 | - | - | - | - | - | - |
| *PDE1C* | 8.94E-25 | - | - | - | - | - | - |
| *VEGFC* | 1.44E-24 | - | - | - | - | - | - |
| *RUNX1* | 3.45E-24 | - | - | - | - | - | - |
| *OSR2* | 3.71E-24 | - | - | - | - | - | - |

<sup>a</sup>Differentially expressed genes. <sup>b</sup>Pluripotent stem cells. <sup>c</sup>False discovery rate. <sup>d</sup>Naked mole-rat. <sup>e</sup>No significant difference. <sup>f</sup>No read detected.

To investigate the functional molecular networks regulating PSCs in each species, we analysed the changes in biological KEGG pathways using GSEA (Fig. 5 and Supplementary Table 6). In general, the activated and suppressed pathways differ among species. Nevertheless, GSEA revealed common activation of translational control pathways, such as ribosome, spliceosome, and nucleocytoplasmic transport. GSEA also revealed frequent activation and suppression of cell adhesion pathways, including focal adhesion, ECM-receptor interaction, and tight junctions in multiple species. Together, our RNA-seq analysis revealed that the comprehensive gene expression of gz-iPSCs had been changed compared to that of gz-fibroblasts by reprogramming, supporting the successful generation of gz-iPSCs.

**Fig. 5.**
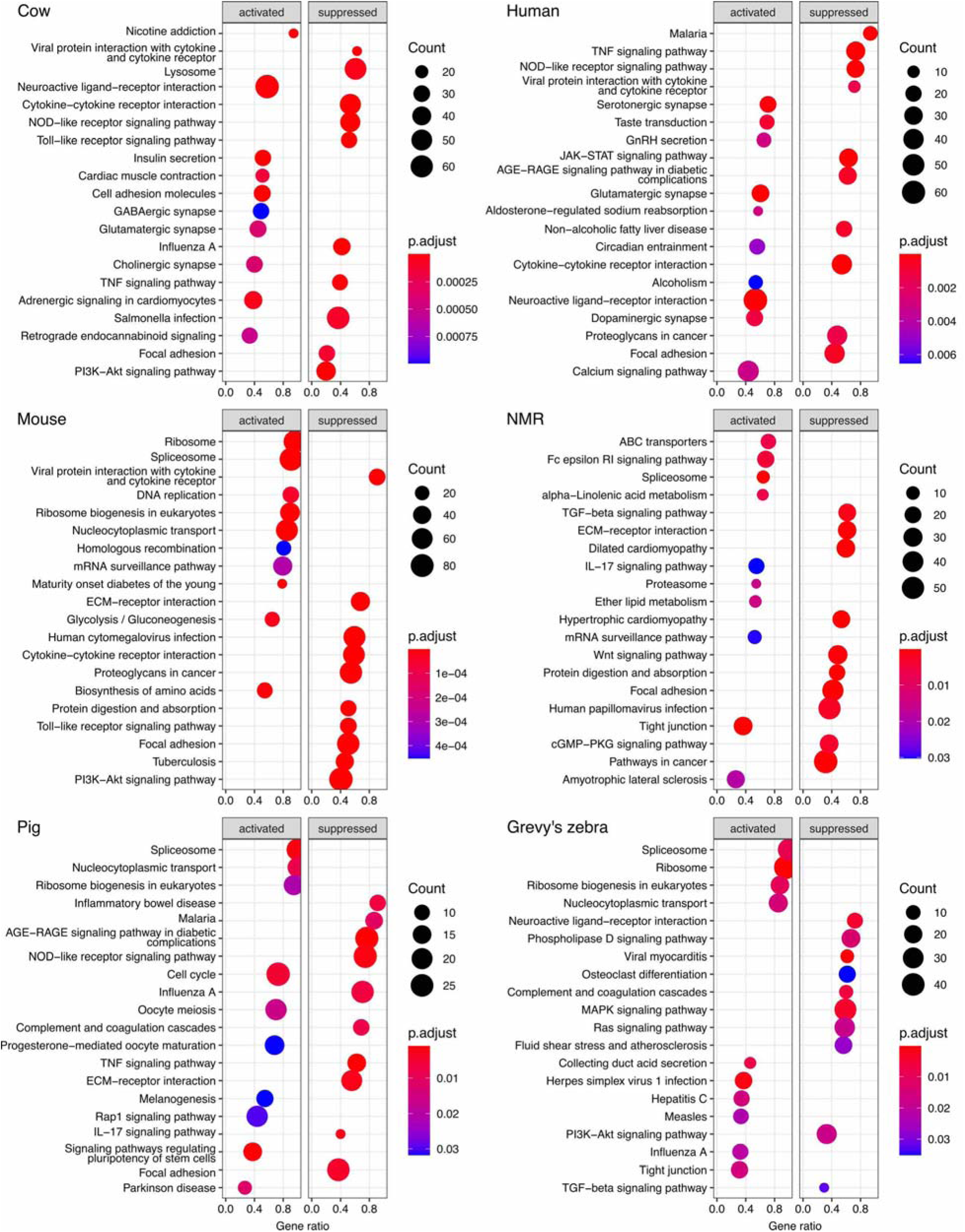
Pathway enrichment of mammalian PSCs. Significantly enriched activated and suppressed KEGG pathways in each species. The horizontal line represents the gene ratio, and the vertical items represent the KEGG terms in order of gene ratio up to 10 pathways each. The depth of the colour represents the adjusted *P-*value, and the size of the circle represents gene counts. PSC, pluripotent stem cells; NMR, naked mole-rat; KEGG, Kyoto Encyclopedia of Genes and Genomes.

## Discussion

In this study, we report the generation of the first iPSCs from an endangered species, Grevy’s zebra. Primary gz-fibroblasts were obtained and successfully reprogrammed into gz-iPSCs. gz-iPSCs generated in this study exhibited PSC characteristics in terms of morphology, expression of pluripotency markers, and differentiation potential into three germ layers.

In light of RNA-seq results, we revealed molecular basis regulating the pluripotency characteristics of the gz-iPSCs. Similar to ESCs, iPSCs can differentiate into three germ layers, maintain high telomerase activity, and exhibit proliferative potential^60^. The observed GO enrichment in embryo development, cell population proliferation, and telomere maintenance indicates that the derived gz-iPSCs have acquired general characteristics of PSCs as also shown with the differentiation experiments into EB and doubling a time similar to that of human iPSCs.

As shown in the Fig. 3, the highly upregulated DEGs were associated with the EMT-MET process that occur during the reprogramming process from fibroblasts to iPSCs^73^. This complicated transition of cell fate is referred to as EMT-MET, where EMT is first activated, followed by its reversed process MET in the early phase of reprogramming^70, 74^. Our findings imply that EMT-MET may have occurred during the reprogramming of gz-iPSCs and that the function and regulation of EMT-MET are conserved in Grevy’s zebra as in humans and mice.

PSCs can exist in multiple pluripotency states, including naïve, primed, and formative state, that is observed in epiblast-like cells (EpiLCs)^75^. Our experimental data indicated that derived gz-iPSCs have primed type morphology and can be cultured in primed conditions for human PSCs, such as mTeSR-1 medium^30^. In our RNA-seq data, we found upregulation of genes indicative of pluripotency and germline cells. In humans, formative EpiLCs express *BST2* at higher levels than the naïve cells^76^ and also express *GATA3*, *VGLL1*, and *CLDN4* uniquely compared to both naïve and primed cells^77^. Our findings, therefore, imply that the pluripotent state of gz-iPSCs may exhibit a mixture of primed and formative states that differs between humans and mice, addressing the complexity of pluripotency state among mammalian PSCs.

Comparative transcriptomics will shed light on the understanding of the conservation and variations in mammalian PSCs, which share many of the criteria for pluripotency but also differ in their characteristics^13, 14, 20^. We observed that hierarchical clustering separates PSCs and fibroblasts, consistent with patterns of gene expression differences between tissues^23^. We found gz-iPSCs nested among other mammalian PSCs, indicating that the global gene expression of the generated gz-iPSCs is similar to those of other mammalian species. gz-iPSCs were most closely clustered with horse ICM, indicating that the expression patterns, at least, partly resolve phylogenetic relationships, as also observed between cows and pigs. The hierarchical pattern, however, did not resolve phylogenetic relationships overall, constructing two major hierarchal groups, one with Grevy’s zebra, horse, and NMR, and the other with cows and pigs, humans, and mice. The independent branch of the mouse, which is the only naive type among the analysed species^42^, may reflect pluripotency status, which shows distinct expression patterns within species^77^. The sister clade includes Grevy’s zebra, horse, and NMR, implying that these species may have unique molecular mechanisms for maintaining their PSCs compared with other widely studied species, addressing the importance of comparative studies across taxonomic groups.

The common changes in gene expression provide insights into the evolutionary conservation of pluripotency mechanisms in mammalian PSCs. We observed the expression of well-known pluripotency-associated genes, including the core pluripotency transcription factor *OCT3/4*^78, 79^, DNA methyltransferase *DNMT3B*^80^, RNA-binding *ESRP1*^81^, and cell adhesion molecules *EPCAM*^82^ and *DSG2*^83^, suggesting that these genes play important roles across taxonomic lineages. The common upregulation of cell adhesion molecules *EPCAM*^82^ and *DSG2*^83^, which are also used as PSC-specific surface markers^82, 83^, indicate that these genes are effective in fluorescence-activated cell sorting for various species. Our GO analysis provided insights into the conservation of biological processes that play important roles in PSCs. For example, we found enrichment of GO terms associated with RNA polymerase II, which regulates transcription in PSCs^84^. As implied in Grevy’s zebra, we also observed enrichment of GO terms associated with the EMT-MET process across mammalian PSCs^70, 74^. Our analysis also revealed that the expression of genes that are not generally associated with PSCs commonly across species, indicating potential functional importance in mammalian PSCs. Together, the common changes in gene expression across taxonomic lineages will provide insights into the principle of molecular mechanisms regulating mammalian PSCs that have been limited to humans and mice.

The pluripotency state and properties of PSCs are maintained by a complex gene network^85^ and require highly orchestrated translation control^86^. We observed common activation of translation control pathways, supporting the important role of the translation process in mammalian PSCs. In addition, we also detected the upregulation of *ESRP1* across mammalian PSCs. *ESRP1* is a splicing regulator and has been shown to play controversial roles in human and mouse PSCs^81, 87–89^. In mice, knockdown of *ESRP1* positively regulates the expression of core pluripotency genes, *OCT3/4*, *SOX2*, and *NANOG*^86^, whereas *ESRP1* promotes the biogenesis of circular RNAs that maintain pluripotency in human PSCs^89^ and enhances human PSC pluripotency^88^. ESRP1 also drives the EMT-MET process by regulating isoform splicing^67, 90^. Collectively, our findings indicate the evolutionary conservation of EMT-MET and associated translation control pathways across species, implying that further understanding of these processes may be key to elucidating the principles of mammalian PSCs.

However, we found variations in activation and suppression in most other biological pathways, as well as a high degree of dispersion in the gene expression patterns of PSCs. These findings may shed light on the species differences in the characteristics of mammalian PSCs and imply the evolution of the unique molecular mechanisms of species for regulating PSCs. The protein-coding sequences of pluripotency-regulating genes have been evolutionarily conserved across mammals^22^. The high variations in gene expression patterns found in this study may suggest that the characteristic variations in mammalian PSCs may be explained by the differences in gene expression.

This report is one of the few cases of the generation of iPSCs from highly endangered species of non-primate taxonomic group^16, 25, 26^. iPSCs derived from endangered species provide biological resources for functionary research and disease investigation. For example, equine piroplasmosis is a tick-borne disease of equids caused by protozoan parasites^91^. When the anthrax outbreak occurred and killed 53 Grevy’s zebras in Kenya, uncertainty with possible adverse effects of vaccination of Grevy’s zebras impeded the immediate application of medical treatment^92^. As safe and effective protocols are especially important for species with declining populations, iPSCs from endangered species could contribute to the development of therapeutic applications^1^.

One prospect of iPSC technology for conservation management is genetic rescue^93^. As the number of individuals declines, it is accompanied with the loss of genetic variation, and the opportunity to preserve viable biomaterials will become increasingly limited. Because the genetic diversity of Grevy’s zebra has been decreasing^28^, cryopreservation of iPSCs from current individuals will contribute to future conservation efforts.

## Conclusions

Grevy’s zebra iPSCs established in this study have advanced our understanding of mammalian PSCs. The effective reprogramming of gz-fibroblasts by human transcription factors supports the plausible conservation of reprogramming mechanisms between humans and equine. The transcriptome of gz-iPSCs allowed us to further characterise the molecular basis of these newly established iPSCs. Comparative transcriptomics with other species has provided new insights into the gene expression patterns of mammalian PSCs, such as evolutionary conservation of the EMT-MET process and translation control. gz-iPSCs will provide resources for future functional studies and conservation management of this endangered species.

## Supporting information

Supplemental informations

## Authors’ contributions

Y. E., M. I-M., and K. K. designed the study and wrote the manuscript; Y. E. performed the experiments and analysed the data; K. H. supervised the cellular experiments; K. O. performed retrovirus reprogramming; H. I. acquired samples and supervised data interpretation; S. T. performed RNA-seq experiments; M. I-M. and K. K. supervised the project. All authors read and approved the final manuscript.

## Supplementary Material

Additional results supporting this article have been uploaded as part of the online electronic supplementary material.

## Acknowledgements

The authors thank N. Yoshida for providing technical support for the experiments; This work was supported by JSPS KAKENHI Grant Numbers 17H03624, 20H00420 (M. I-M.) and 17H02083 (K.K.), the Kyoto University Supporting Program for Interaction-based Initiative Team Studies (SPIRITS) (M. I-M), and the Environment Research and Technology Development Fund (FPMEERF20214001) of the Environmental Restoration and Conservation Agency of Japan (M. I-M).

## Data availability

RNA-seq data have been deposited under BioProject PRJNA748892 and GEO GSE180619.

Correspondence and requests for Grevy’s zebra cells should be addressed to K.K.

