## Supplemental informations for "Generation of iPSCs from endangered Grevy’s zebra and comparative transcriptomic analysis of mammalian PSCs"

*Corresponding authors

**Supplementary methods**

**Cell culture for human iPSCs**

The human iPSCs (253G1)^1^ were cultured in mTeSR-1 (Stemcell Technologies, Vancouver, Canada)^2^ on Matrigel (Corning, Corning, NY, United States)-coated dishes. Human iPSCs were passaged using TrypLE Express (Thermo Fisher Scientific) with the addition of Rho-associated coiled-coil forming kinase (ROCK) inhibitor Y-27632 (Fujifilm Wako Pure Chemical Corporation, Osaka, Japan) at 10 µM after passaging for 24 h. The cells were cultured in a humidified incubator at 37 °C with 5% (v/v) CO_2_.

**RT-PCR**

Reverse transcription-polymerase chain reaction (RT-PCR) was performed using PrimeScript RT Master Mix (Takara Bio, Shiga, Japan). The cycling conditions for RT-PCR were as follows: 94 °C, 1 min, followed by 35 amplification cycles (94 °C, 30 s; 60 °C, 30 s; 72 °C, 30 s). RT-PCR products were analyzed on 2 % agarose gels (50 V for 45 min).

**Supplementary Figures**

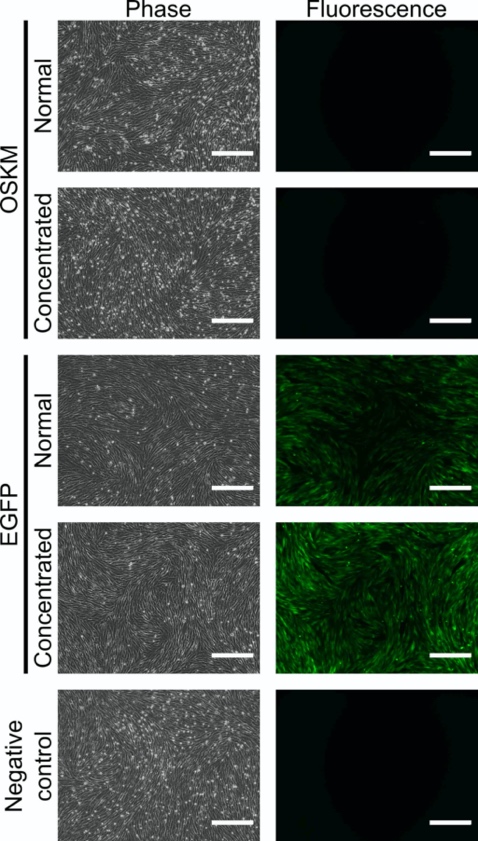

**Supplementary Figure 1 | Morphology of Grevy’s zebra fibroblasts after transduction.**

Grevy’s zebra fibroblasts were transduced with either normal or concentrated retroviral vectors. Cellular reprogramming was conducted with retroviral vectors for *OCT3/4*, *SOX2*, *KLF4*, and *c-MYC* (OSKM), and transfection efficiency was observed with EGFP expression. The negative control represents Grevy’s zebra fibroblasts cultured in a normal fibroblast medium without viral transduction. The cells were photographed at 3 days after transduction. The scale bar represents 300 μm.

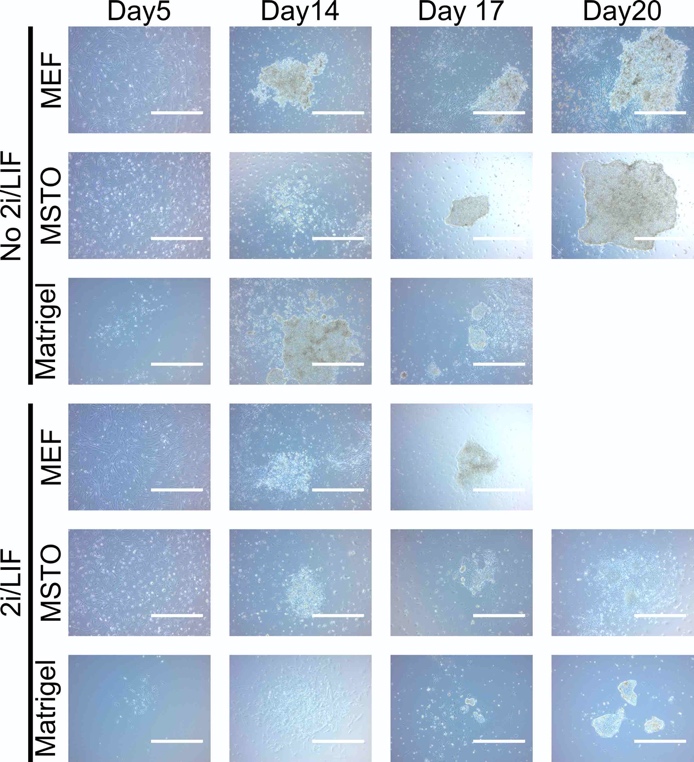

**Supplementary Figure 2 | Formation of iPSC-like colonies with different culture conditions at reprogramming.**

The transduced Grevy’s zebra fibroblasts were reseeded on day 3 after transduction and cultured with different culture conditions. Cells transduced with the concentrated vector are shown here. Photographs for without 2i/LIF on Matrigel and with 2i/LIF on MEF were not taken because the colonies were selected on day 17. The scale bar represents 1000 μm. No PSC-like colony was observed in without 2i/LIF on MEF. MEF, mouse embryonic fibroblasts; MSTO, mouse SNL-STO; 2i/LIF, CHIR99021, PD0325901 (2i), and leukaemia inhibitory factor (LIF).

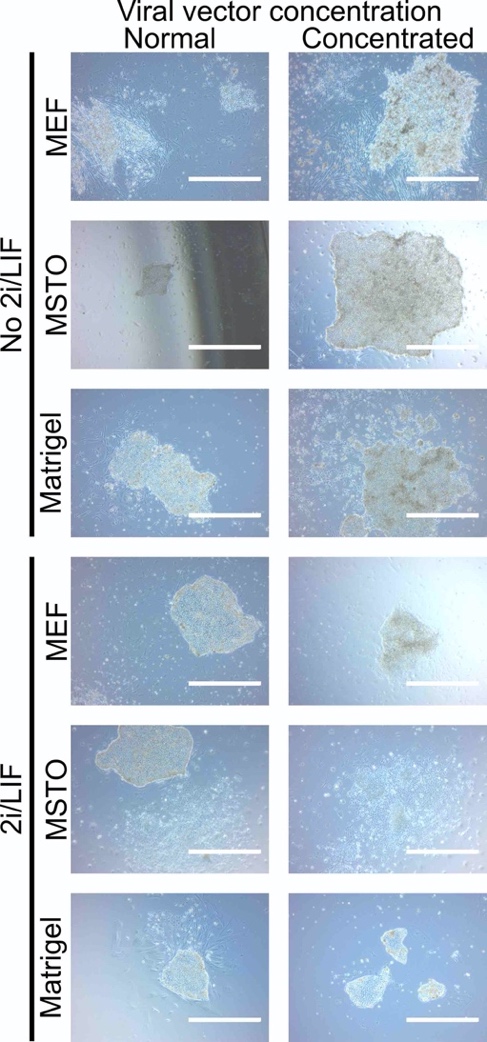

**Supplementary Figure 3 | Morphology of PSC-like colonies with different culture conditions at reprogramming.**

Colonies with PSC-like morphology appeared in all tested conditions. No PSC-like colony was observed in conditions without 2i/LIF on MEF. The morphology of the derived colonies was similar across conditions independence of 2i/LIF. Photographs were taken from days 14–20. The scale bar represents 1000 μm. PSC, pluripotent stem cells; MEF, mouse embryonic fibroblasts; MSTO, mouse SNL-STO; 2i/LIF, CHIR99021, PD0325901 (2i), and leukaemia inhibitory factor (LIF).

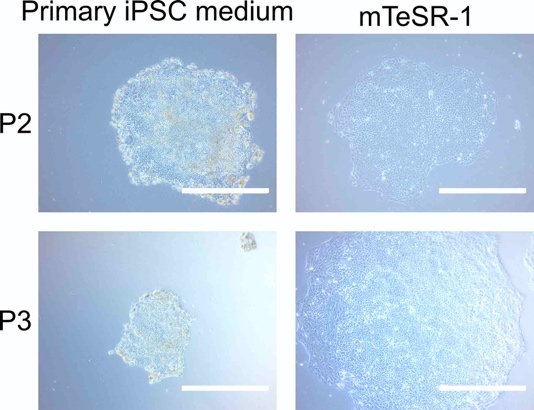

**Supplementary Figure 4 | Growth of derived gz-iPSCs under primary iPSC medium and mTeSR-1.**

The putative gz-iPSCs exhibited heterogeneous morphology and stopped growing after three to five generations in the primary iPSC medium. In contrast, the cells formed homogenous and dense colonies and kept growing in mTeSR-1. Photographs were taken on day 8 after reseeding for passage 2 and on day 4 for passage 3. The scale bar represents 1,000 μm. gz-iPSCs, Grevy’s zebra induced pluripotent stem cells.

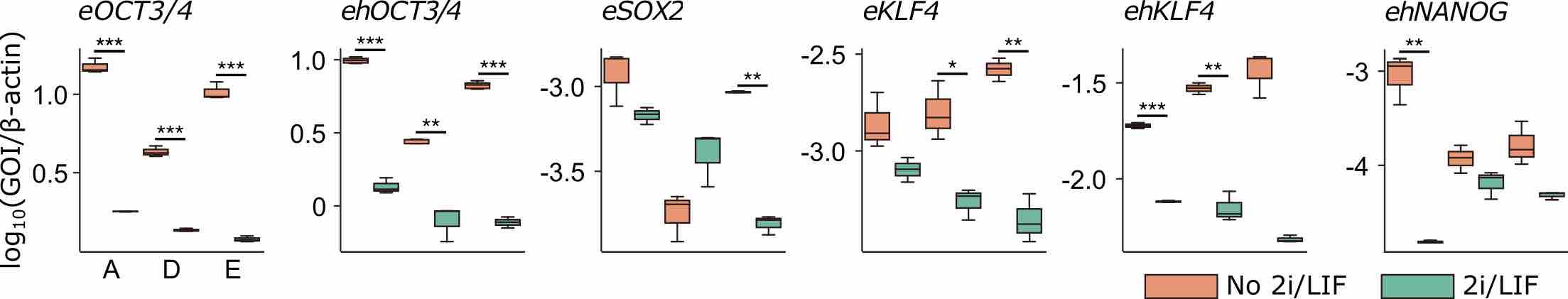

**Supplementary Figure 5 | 2i/LIF treatment of gz-iPSCs.**

Comparison of pluripotency marker expressions between gz-iPSCs cultured with or without 2i/LIF using qRT-PCR. A, D, and E represents independent clones of gz-iPSCs at passage 25 without 2i/LIF and 29, 32, and 29 with 2i/LIF, respectively. Three independent experiments were performed. Centre lines indicate median and box limits indicate upper and lower quartiles. Upper whisker = min(max(*x*), Q_3 + 1.5 × IQR), lower whisker = max(min(*x*), Q_1 - 1.5 × IQR). Statistical analyses were performed using the Welch two-sample t-test (* *P* < 0.05, ** *P* < 0.01, and *** *P* < 0.001). *eOCT3/4*, A with 2i/LIF-A without 2i/LIF, df = 2.0054, *P* = 0.0008597, D with 2i/LIF-D without 2i/LIF, df = 2.325, *P* = 0.0007303, E with 2i/LIF-E without 2i/LIF, df = 2.4499, *P* = 0.0004037; *ehOCT3/4*, A with 2i/LIF-A without 2i/LIF, df = 2.8042, *P* = 0.0002151, D with 2i/LIF-D without 2i/LIF, df = 4.9878, *P* = 0.003064, E with 2i/LIF-E without 2i/LIF, df = 4.6007, *P* = 1.14 × 10^-6^; *eSOX2*, A with 2i/LIF-A without 2i/LIF, df = 2.3798, *P* = 0.1124, D with 2i/LIF-D without 2i/LIF, df = 3.91, *P* = 0.05003, E with 2i/LIF-E without 2i/LIF, df = 2.0343, *P* = 0.001617; *eKLF4*, A with 2i/LIF-A without 2i/LIF, df = 2.7379, *P* = 0.08896, D with 2i/LIF-D without 2i/LIF, df = 3.0565, *P* = 0.01805, E with 2i/LIF-E without 2i/LIF, df = 2.8845, *P* = 0.002741; *ehKLF4*, A with 2i/LIF-A without 2i/LIF, df = 2.3701, *P* = 0.0001695, D with 2i/LIF-D without 2i/LIF, df = 2.5856, *P* = 0.002024, E with 2i/LIF-E without 2i/LIF, df = 2.0946, *P* = 0.005261; *ehNANOG*, A with 2i/LIF-A without 2i/LIF, df = 2.0127, *P* = 0.007246, D with 2i/LIF-D without 2i/LIF, df = 3.9997, *P* = 0.09927, E with 2i/LIF-E without 2i/LIF, df = 2.1248, *P* = 0.05338. In all statistical tests, sample size is n = 3, except *ehOCT3/4* with A, D, and E without 2i/LIF, n = 6. gz-iPSCs; Grevy’s zebra induced pluripotent stem cells; 2i/LIF, CHIR99021, PD0325901 (2i), and leukaemia inhibitory factor (LIF); df, degrees of freedom; *e* and *h* represent genes of equine and human, respectively. gz-iPSCs, Grevy’s zebra induced pluripotent stem cells.

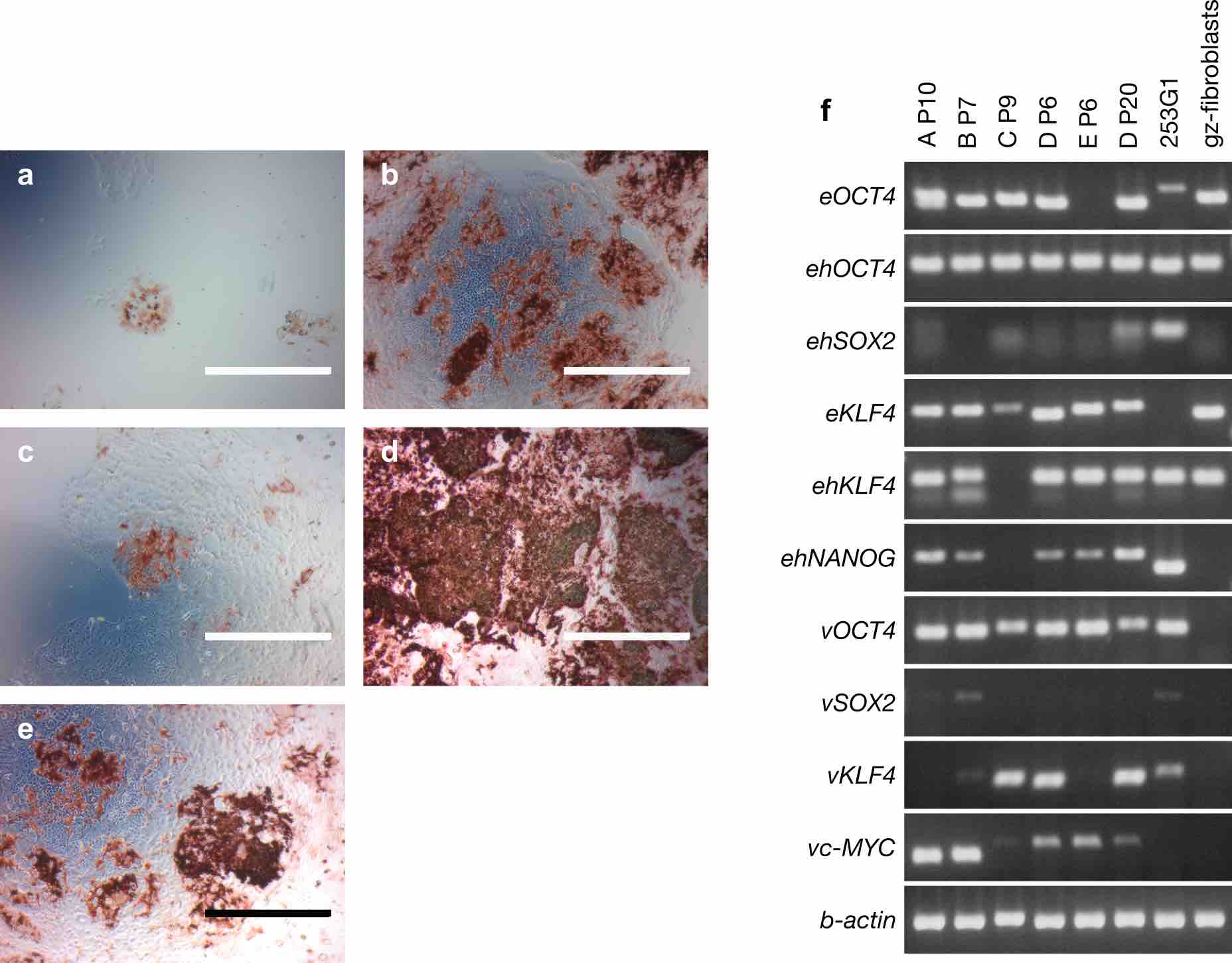

**Supplementary Figure 6 | Preliminary pluripotency experiments for gz-iPSC colony selection.**

**a–e,** Alkali phosphatase (AP) activity of gz-iPSC clone A P6, B P8, C P10, D P8, and E P10. Clone B, D, and E show high AP activity while clone A and C show weak AP activity. The scale bar represents 1,000 μm. **f**, RT-PCR for testing primer reaction and colony selection. To test the primer reaction and select primary colonies, we performed RT-PCR. A, B, C, D, and E represents derived gz-iPSC clones, and P represents passage number. D P20 was used to compare gene expression at later passages. Human iPSC (253G1) was used as pluripotent stem cell control and Grevy’s zebra fibroblasts were used as fibroblast control. We observed that all the primers reacted as expected. Although the equine-specific primer *eOCT4* reacted with Grevy’s zebra and human sample, the human PCR product was longer and lower in intensity compared with those of Grevy’s zebra. Considering that the PCR products with *ehOCT4* primer, which reacts with both human and *Equus* genes, were stable in length and intensity across samples, Grevy’s zebra samples, which expressed *OCT4* and *eOCT4* primer more specifically reacted with the host gene rather than the viral gene. Clone D P20 showed relatively higher expressions with *ehSOX2* and *ehNANOG* compared with samples at earlier generations, suggesting an increase in expression level at later generations. In selecting primary clones for later analysis, we excluded clone C, which showed low expression of *eKLF4* and no expression of *ehKLF4* and *ehNANOG*. We also excluded clone B, which expressed all the four viral genes. Clone A shows relatively weak AP activity yet exhibits expressions of all pluripotency markers, including the highest expression of *ehNANOG*, and silencing of *vSOX2* and *vKLF4*. Therefore, we continued culturing the selected clones A, D, and E and used them for later analysis. *e*, equine; *h*, human; *v*, viral.

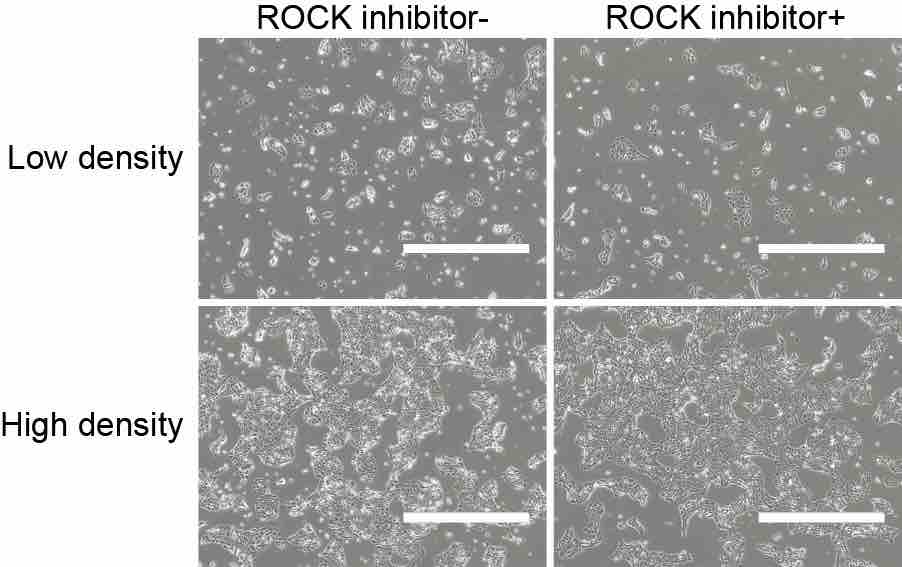

**Supplementary Figure 7 | ROCK inhibitor treatment of gz-iPSCs.**

gz-iPSCs were treated with Rho-associated coiled-coil forming kinase (ROCK) inhibitor Y-27632 for 24 h before and after passaging. TrypLE Express was used to dissociate the cells. The cells attached to the dish without ROCK inhibitor, but we observed more floating cells and obscure edges of colonies. ROCK inhibitor improved cell attachment to the dish, and the colonies exhibited distinct edges. gz-iPSC clone D at passage 31 was used. Photographs were taken on day 1 after passaging at low- and high-density areas in a dish. The scale bar represents 1,000 μm. gz-iPSCs, Grevy’s zebra induced pluripotent stem cells.

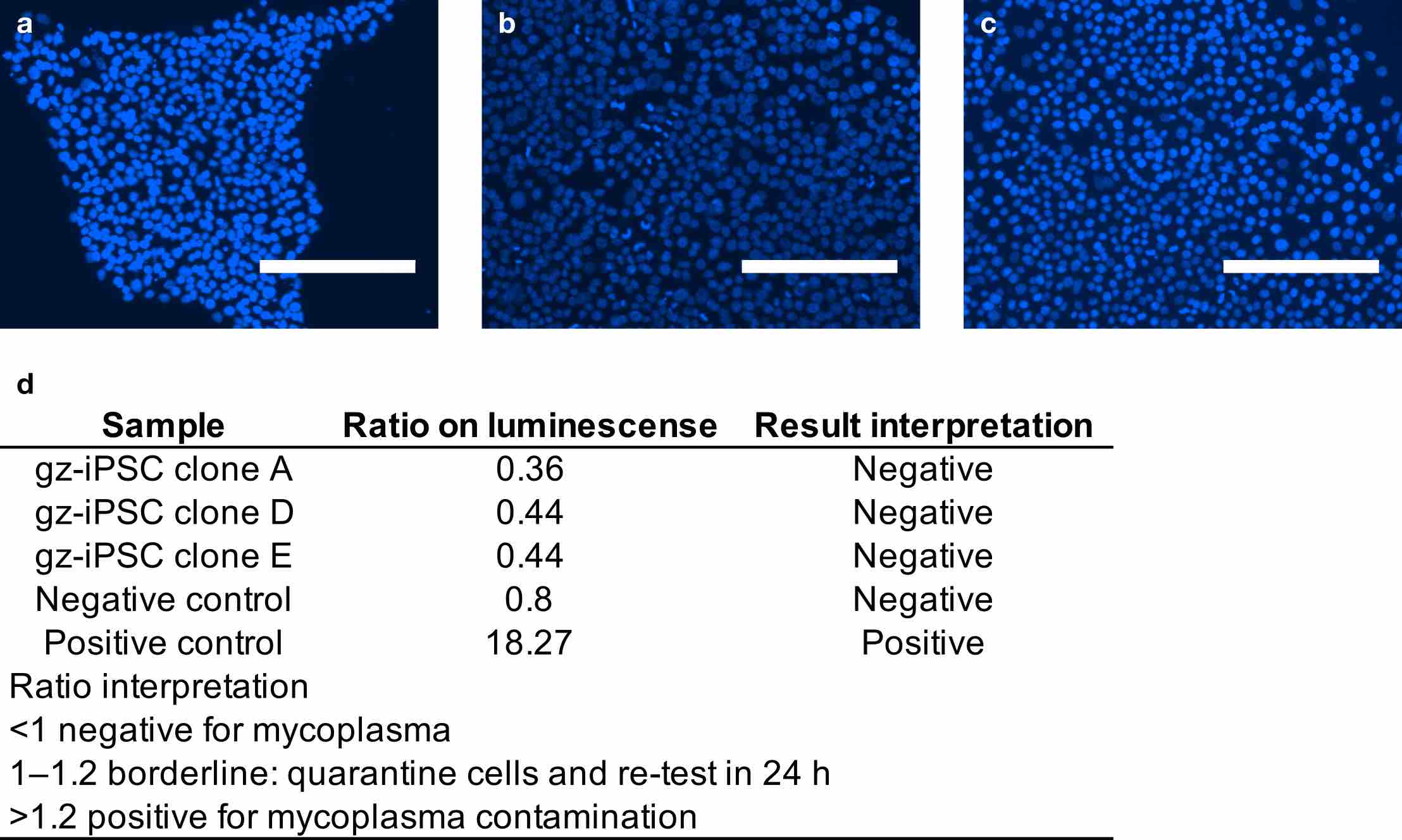

**Supplementary Figure 8 | Mycoplasma detection of gz-iPSCs.**

**a–c,** gz-iPSC clones A, D, and E at passage 32 using nuclear staining with DAPI. The scale bar represents 200 μm. **d,** Mycoalert results on gz-iPSC clones. gz-iPSCs, Grevy’s zebra induced pluripotent stem cells; DAPI, 4ʹ,6-diamidino-2-phenylindole.

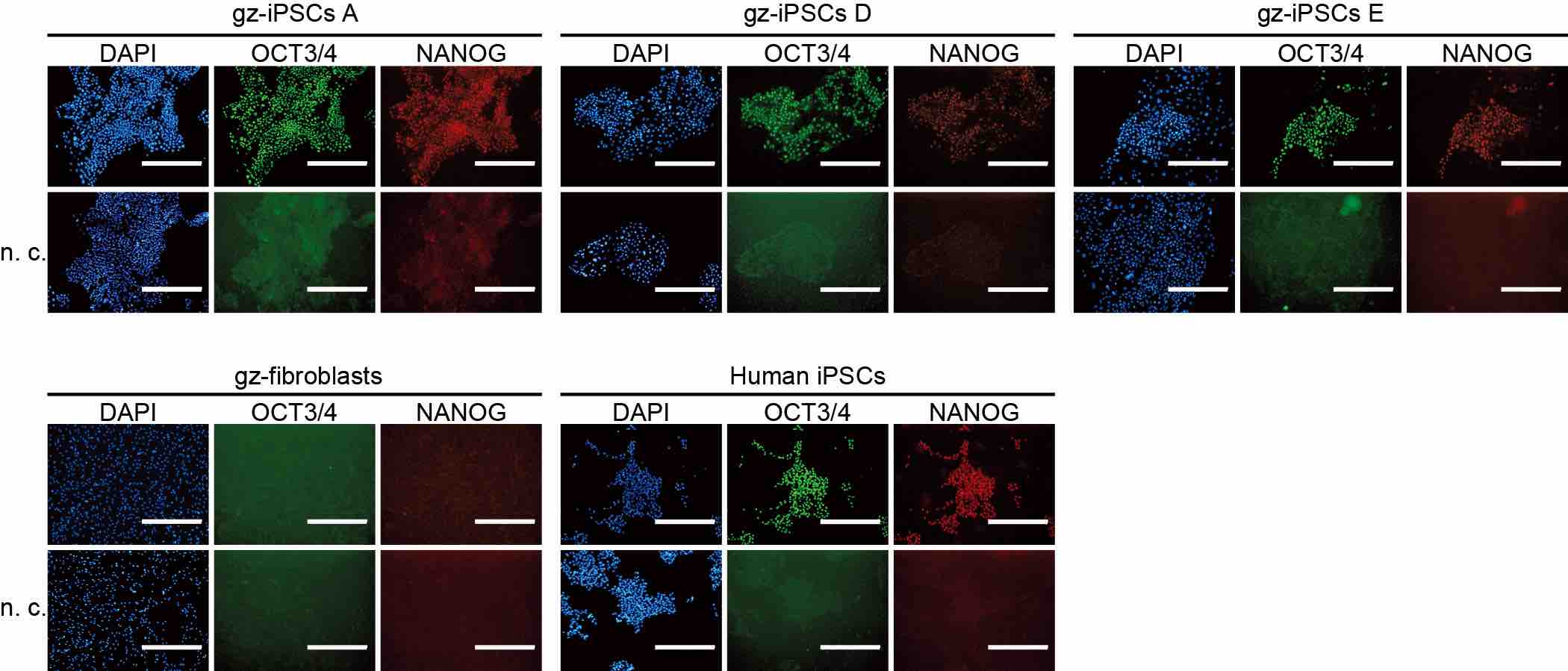

**Supplementary Figure 9 | Immunofluorescence for the pluripotency markers.**

Immunostaining for OCT3/4 and NANOG are shown for three independent clones of gz-iPSCs (A, D, and E at passage 17), gz-fibroblasts, and human iPSCs (253G1). Nuclei are stained with DAPI, 4ʹ,6-diamidino-2-phenylindole. The fluorescent expression in the human iPSCs confirms the successful staining with antibodies. No fluorescent expression in the negative controls supports that secondary antibodies reacted with primari antibodies. No fluorescent expression in the fibroblasts supports that expression of pluripotency markers was increased in the generated gz-iPSCs. The scale bar represents 400 μm. gz-iPSCs, Grevy’s zebra induced pluripotent stem cells; n.c., negative controls.

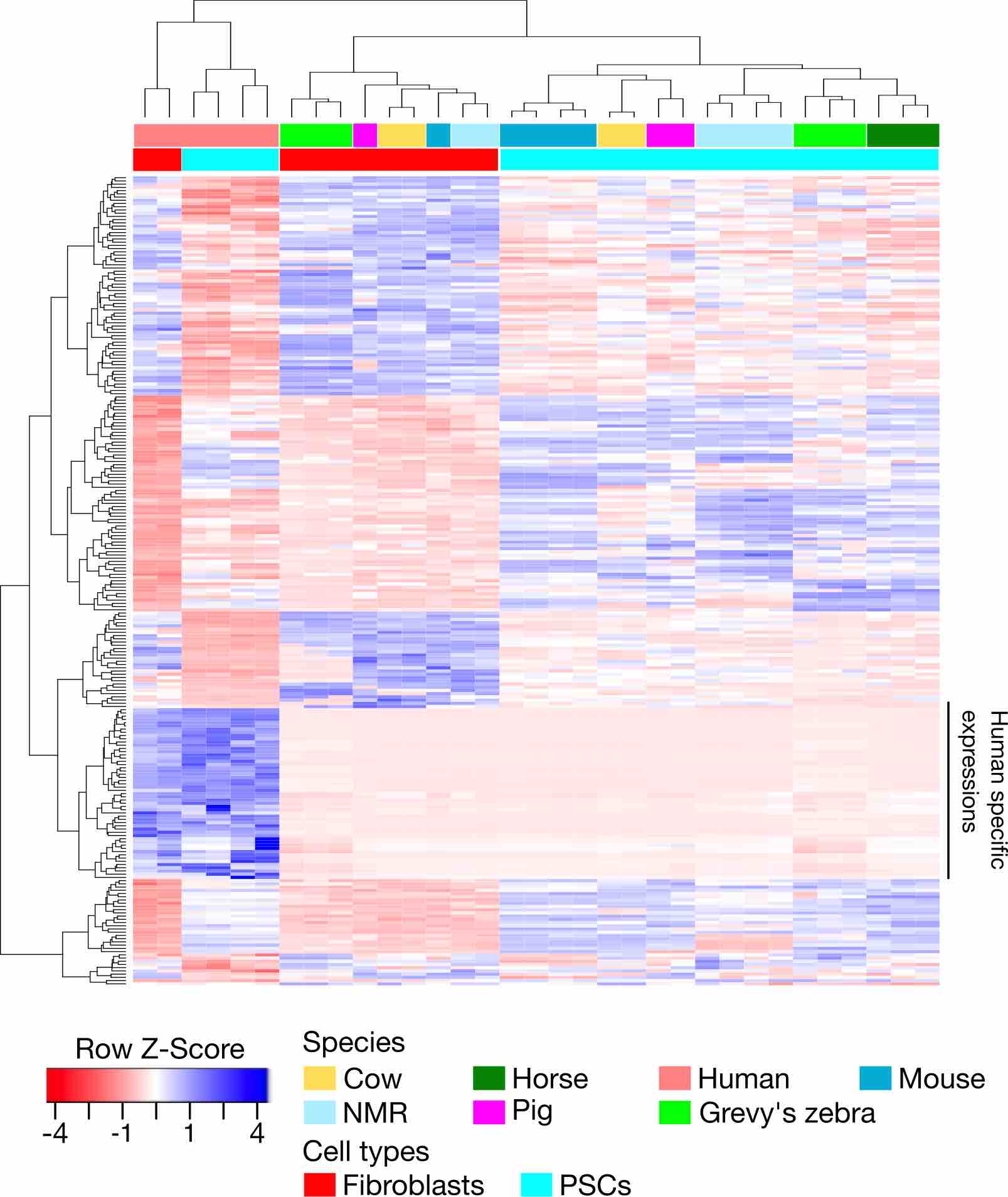

**Supplementary Figure 10 | A heat map with hierarchical clustering of DEGs including genes expressed only in humans.**

DEGs between PSCs and fibroblasts were estimated across species. The colour bars at the top indicate species, and those at the bottom indicate cell types. The samples were clustered by cell types, except humans. Human samples, both PSCs and fibroblasts, show unique expression of genes that are not observed in other species, putatively caused by annotation bias. To avoid this problem, we excluded genes with which RNA sequencing read was detected only in human samples. PSCs, pluripotent stem cells; DEGs, differentially expressed genes; NMR, naked mole-rat.
